# NAK associated protein 1/NAP1 is required for mitosis and cytokinesis by activating TBK1

**DOI:** 10.1101/2022.03.09.483647

**Authors:** Swagatika Paul, Shireen A. Sarraf, Ki Hong Nam, Leila Zavar, Sahitya Ranjan Biswas, Lauren E. Fritsch, Nicole DeFoor, Tomer M. Yaron, Jared L. Johnson, Emily M. Huntsman, Lewis C. Cantley, Alban Ordureau, Alicia M. Pickrell

## Abstract

Subcellular location and activation of Tank Binding Kinase 1 (TBK1) govern precise progression through mitosis. Either loss of activated TBK1 or its sequestration from the centrosomes causes error in mitosis and growth defects. Yet, what regulates its recruitment and activation on the centrosomes is unknown. We identified that NAK Associated Protein 1 (NAP1) is essential for mitosis which binds to TBK1 on the centrosomes to activate it. Loss of NAP1 causes several mitotic and cytokinetic defects due to inactivation of TBK1. Our quantitative phosphoproteomics identified numerous TBK1 substrates that are not only confined to the centrosomes but also are associated with microtubules. Substrate motifs analysis indicates that TBK1 acts upstream of other essential cell cycle kinases like Aurora and PAK kinases. We also identified NAP1 as a TBK1 substrate at S318 promoting its degradation by ubiquitin proteasomal system acting as a negative regulatory step. These data uncover an important distinct function for the NAP1-TBK1 complex during cell division.

## Introduction

Successful cell division is dependent on the precise and timely transition between different cell cycle phases, which is regulated by dynamic changes in protein phosphorylation. Thus, protein kinases play a vital role in orchestrating almost every step of cell division including DNA replication, centrosome maturation, chromatin condensation, spindle assembly formation, sister chromatid segregation, and cytokinesis [1-4]. Entry into mitosis is marked by the highest incidences of protein phosphorylation and kinase activity [5, 6]. Therefore, impaired or aberrant kinase activity often leads to errors in all these cell cycle events which consequently become the underlying cause for developmental defects [7, 8] or abnormal cell proliferation leading to cancer [9, 10]. Tank Binding Kinase 1 (TBK1) is one such kinase which is known to be activated on the centrosomes during mitosis [11, 12] and is also often overexpressed in certain cancer types [13-15]. Genetic loss of TBK1 leads to embryonic lethality in mice [16], and its loss results in mitotic defects in cancer cell lines [11, 12, 17]. Interestingly, sequestration of activated TBK1 away from the centrosomes also disrupts mitosis in neural epithelial stem cells and radial glia during Zika virus infection [18]. Our previous work has shown that sequestration of TBK1 to the mitochondria during mitophagy, the selective degradation of mitochondria, also blocks mitosis due to the unavailability of activated TBK1 on the centrosomes [12]. Thus, both proper activation and subcellular localization of TBK1 are essential for mitotic progression. Yet, the upstream regulation of TBK1 during mitosis is unknown, and we do not completely understand the function of activated TBK1 on the centrosomes.

Activation of TBK1 depends on its binding to an adaptor protein which induces a conformational change leading to trans-autophosphorylation on serine 172 of the kinase domain of TBK1 [27, 36, 37, 39]. Interaction with the adaptor protein not only activates the TBK1 kinase domain but may drive its subcellular localization to different organelles to regulate distinct signaling pathways [19-21]. From extensive studies examining its regulation during innate immune signaling, autophagy, and mitophagy, we know that TBK1 has multiple binding partners/adaptors for each of these cellular processes. While TANK, SINTBAD, NAP1/AZI2, optineurin (OPTN), and STING [20, 22-29] are the major TBK1 adaptors during innate immune signaling, OPTN, TAX1BP1, and NDP52 [21, 30-32] can activate TBK1 during mitophagy. However, the adaptor or adaptors required for TBK1 activation and recruitment during mitosis are unknown.

Along with activation, localization of TBK1 on the centrosomes is also essential for mitosis [12, 18]. This localization brings activated TBK1 in the proximity of microtubules as centrosomes are the microtubule organizing center. Past studies have identified a few of the TBK1 substrates on the centrosomes [11, 17, 33]. Whether TBK1 functions only to phosphorylate centrosomal proteins during mitosis or also regulates microtubule binding proteins remains to be determined. The complete landscape of the proteins targeted by TBK1 during mitosis remains unclear. Identifying all of the mitotic substrates would offer mechanistic insight into pathways modulated by TBK1 to ensure proper chromosomal segregation.

We show that NAP1/AZI2 is necessary for TBK1 activation during mitosis, whose function has only been described in innate immunity to trigger Type I interferon or NF-κB signaling [26, 34, 35]. We discovered NAP1 to be a centrosomal protein which regulates proper cell division by binding and activating TBK1 during mitosis. Loss of either NAP1 or TBK1 result in the accumulation of binucleated and multinucleated cells, due to the several mitotic and cytokinetic defects observed across several cell lines. We also describe a new function for both proteins, as our data suggests that they are also implicated in cytokinesis. Interestingly, NAP1 levels during mitosis are tightly regulated by TBK1. Activated TBK1 phosphorylates NAP1 on serine 318 flagging it for ubiquitin proteasomal degradation (UPS). Through unbiased quantitative phosphoproteomics analysis during mitosis, we also uncovered unidentified TBK1 substrates, which implicate its upstream effects on other cell cycle kinases such as Aurora A and Aurora B. This NAP1-TBK1 signaling axis during mitosis is unique to its function in innate immunity.

## Results

### NAP1/AZI2 is required for TBK1 activation during mitosis

Activation of TBK1 is reliant upon its binding to adaptor proteins, which initiates higher order oligomerization of the TBK1-adaptor complex, leading to trans-autophosphorylation at serine 172 (p-TBK1) [36-38]. Previous identification of these adaptor proteins for TBK1 activation in other cellular contexts display overlap and redundancy. For example, during mitophagy, TAX1BP1, optineurin, and NDP52 have been found to be necessary for TBK1 activation [21, 30-32]. Optineurin and NDP52 are also the bound adaptor for TBK1 during certain innate immune stimuli [36, 39-41]. Despite these observations, the adaptor or adaptors required for TBK1 during miosis are unknown. Therefore, we screened known TBK1 adaptors to determine whether any of these proteins were responsible for its activation during mitosis.

First, we investigated TBK1 adaptors restricted to innate immune signaling, TANK, SINTBAD, and NAP1, by generating cell lines in which these proteins were stably knocked down (KD) at the mRNA and protein level in HeLa cells (**Figure 1A-F**). Reductions in TANK and SINTBAD did not alter TBK1 activation during mitosis (**Figure 1D-E**). However, NAP1 KD cells displayed a reduction of p-TBK1 during mitosis, indicating NAP1 could be required for TBK1 activation (**Figure 1F**). We also assessed p-TBK1 levels in an NDP52/OPTN double KO (DKO) HeLa cell line as both adaptor proteins have been implicated in TBK1 activation during mitophagy, xenophagy, and innate immunity [21, 30-32, 40, 42-46], but found no difference between p-TBK1 levels when synchronized (**Figure 1G**). This was also true for a cell line lacking five autophagy related adaptors that TBK1 either binds or phosphorylates: NBR1, TAX1BP1, OPTN, p62, and NDP52 (**Figure 1H**).

**Fig. 1.**
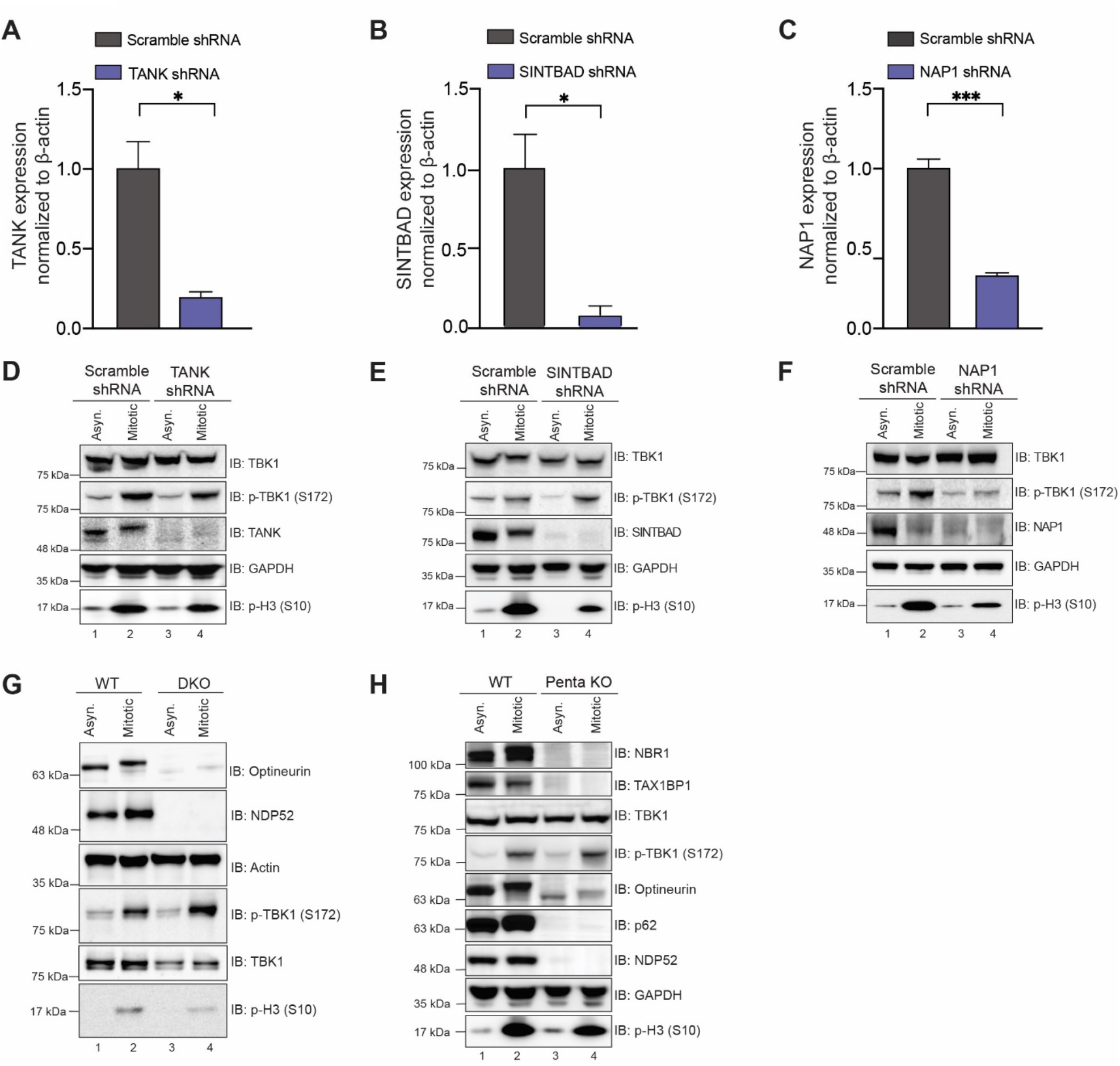
Other known TBK1 adaptors except NAP1 are not required for TBK1 activation during mitosis. (**A-C)** qRT-PCR showing relative expression of TANK1 **(A)**, SINTBAD **(B)** and NAP1 **(C)** mRNA levels normalized to Δ-actin in HeLa cells stably expressing shRNAs. n=3 independent experiments. Error bars ±SD. **(D-F)** Western blot analysis of p-TBK1 in asynchronous and synchronized mitotic cells from scramble control, and cell lines stably expressing TANK **(D)**, SINTBAD **(E)**, and NAP1 **(F)** shRNAs, respectively. Nocodazole was used for cell synchronization. **(G)** Western blot analysis of p-TBK1 levels during mitosis in NDP52/optineurin DKO and parental HeLa cells in asynchronous and mitotic conditions. Nocodazole was used for cell synchronization **(H)** Western blot analysis of p-TBK1 levels during mitosis in HeLa and Penta KO HeLa cells lacking NBR1, TAX1BP1, optineurin, NDP52, p62 in asynchronous and mitotic cells. Nocodazole was used for cell synchronization. Unpaired Student’s t-test was performed for all statistical analysis. * p < .05, *** p <.001.

To confirm that TBK1 activation during mitosis was dependent on NAP1, we generated two independent NAP1 CRISPR knockout (KO) clones targeting exon 4 (**Figure 2A**). Both NAP1 KO HeLa clones displayed decreased p-TBK1 levels during mitosis (**Figure 2A**). To confirm this result, we performed immunocytochemical experiments to detect p-TBK1 staining on centrosomes, which was significantly reduced in NAP1 KO cells (**Figure 2B-C**). This result was specific to NAP1 because p-TBK1 levels were restored during mitosis upon stable reintroduction of the protein (**Figure 2D**).

**Fig. 2.**
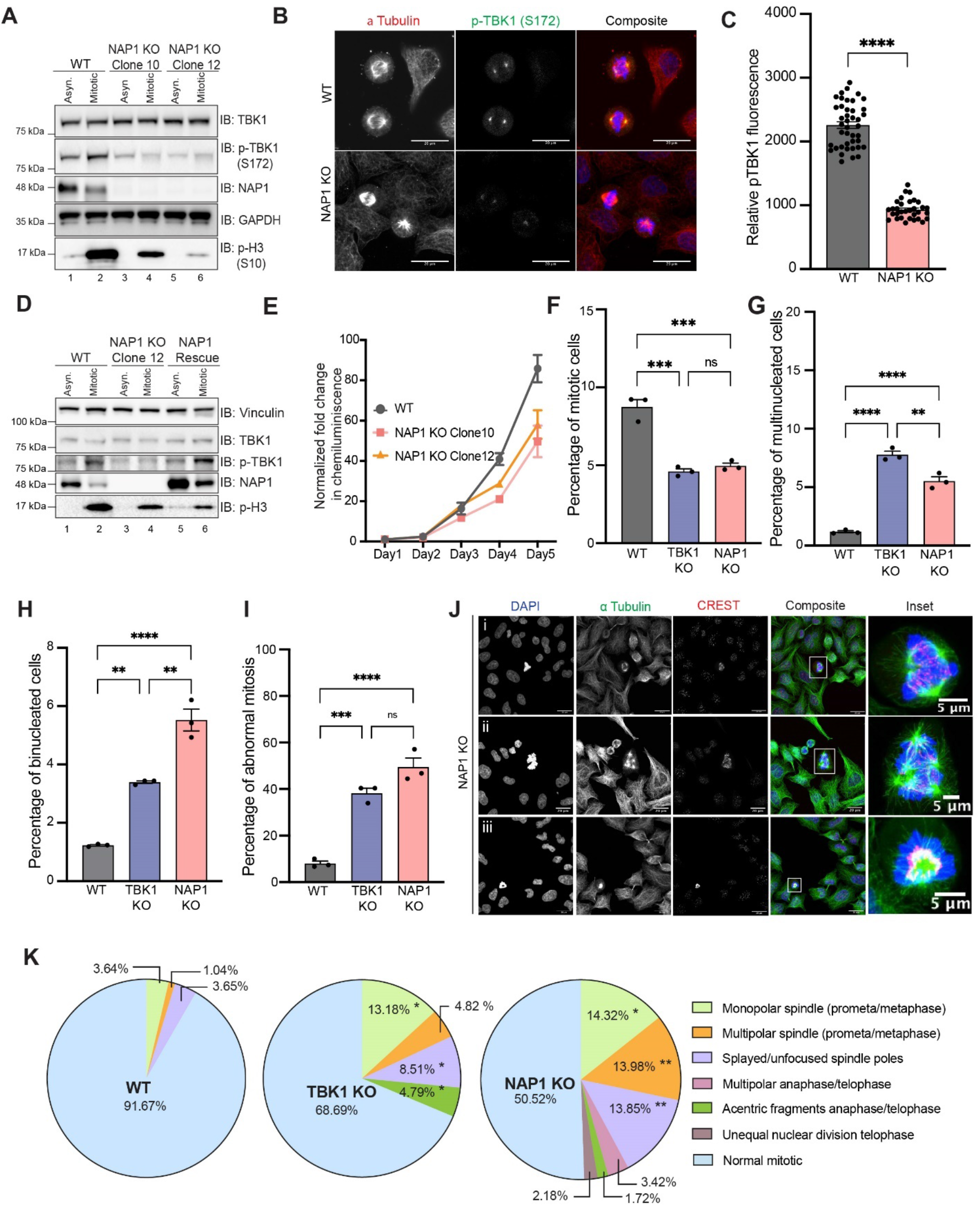
NAP1 KO cells have mitotic defects similar to those lacking TBK1. **(A)** Western blot analysis of p-TBK1 in asynchronous and mitotic cells from WT HeLa, NAP1 KO clone 10, and NAP1 KO clone 12. Nocodazole was used for cell synchronization at mitosis. **(B)** Representative confocal images of p-TBK1 expression on mitotic centrosomes from WT HeLa and NAP1 KO cells. DAPI (blue) was used as a nuclear counterstain (on composite image), α-tubulin for cytoskeleton staining (red), and p-TBK1 S172 conjugated 488 (green). Scale bar - 20 μm. **(C)** Relative intensity of p-TBK1 on centrosomes of mitotic cells from WT HeLa and NAP1 KO cells. 40-50 mitotic cells per group were quantified from 2 biological replicates. Error bars indicate ±SEM. **(D)** Western blot analysis of p-TBK1 in asynchronous and synchronized mitotic cells from HeLa, NAP1 KO clone 12, and the stable NAP1 rescue line. RO-3306 was used for cell synchronization prior to mitotic release. **(E)** Growth curve with normalized luminescence for WT HeLa cells, NAP1 KO clone 10 and NAP1 KO clone 12. Error bars indicate ±SD for technical replicates. (**F-I)** Mitotic (F), multinucleated (G), binucleated (H), and abnormal mitotic (I) percentage cell counts from an asynchronous population of WT HeLa, TBK1 KO, and NAP1 KO cells. Error bars indicate ±SEM; n=3 independent experiments. For mitotic index, multinucleated, and binucleated cell counts, random fields of view were captured sampling approximately 1000 cells per biological replicate from each genotype. For abnormal mitotic cell counts, random fields of view were captured to sample approximately 50 mitotic cells per biological replicate from each genotype. **(J)** Representative confocal images of NAP1 KO mitotic defects: (i) multipolar metaphase, (ii) multipolar prometaphase, and (iii) monopolar prometaphase/metaphase cells. DAPI (blue) was used as a nuclear counterstain, α-tubulin for cytoskeleton staining (green), and CREST for kinetochore staining (red). Scale bar, 20 μm, insets, 5μm. **(K)** Pie charts representing the percentage of different types of mitotic defects found in WT HeLa, TBK1 KO, and NAP1 KO cells. At least 50 mitotic cells per biological replicate from each genotype were analyzed. n=3 independent experiments. One way ANOVA was performed for all statistical analysis. * p < .05, ** p < .01, *** p <.001, **** p <.0001, ns = not significant.

### NAP1 KO cells have mitotic and cytokinetic defects like those lacking TBK1

Previously, we and others have shown that loss of TBK1 led to slowed cell growth, decreased number of mitotic cells in asynchronous conditions, and an increased prevalence of multinucleated cells [11, 12, 18, 33]. However, a more thorough analysis of the defects in cell division that led to these observations have not been performed. Therefore, we characterized cell division defects in NAP1 and TBK1 KO HeLa cells to compare whether these lines phenocopied each other. In asynchronous conditions, NAP1 KO cells exhibited slower growth rates, fewer number of mitotic cells, an increase in the number of bi- and multinucleated cells, and an increased number of cells displaying abnormal mitotic division (**Figure 2E-I**). We further characterized the types of abnormal mitotic defects between these KO lines. Both genotypes had a significant prevalence of monopolar spindles and splayed/unfocused spindle poles, but TBK1 KO had a significantly higher percentage of acentric fragments, while NAP1 KO cells had a higher percentage of cells with multipolar spindles (**Figure 2J-K**).

To further characterize how NAP1 and TBK1 regulate the progression of cell division, we also evaluated if cytokinetic defects were present. Both KO lines had a significantly higher percentage of cytokinetic cells in asynchronous conditions that display a higher percentage of cytokinetic defects (**Figure S1A-B**). Both KO lines exhibited unequal cytokinesis, as well as multipolar cytokinesis at a higher percentage than the parental line (**Figure S1C-D**).

We attempted to use the non-transformed near diploid RPE-1 and DLD-1 cell lines to confirm our findings in HeLa cells but were unable to generate either knockout or stable shRNA-mediated knockdown of NAP1 in both cell lines due to excessive cell death. Using a transient viral mediated transduction of NAP1 shRNA over 36 hours, we could generate a transient reduction of NAP1 (**Figure 3A**). These transient NAP1 KD in DLD-1 cells had reduced p-TBK1 levels during mitosis (**Figure 3A**). From the mitotic analysis on transient NAP1 KD DLD-1 cells, we observed a decrease in the mitotic index and increased percentage of multinucleated cells in transient (**Figure 3B-E**). However, there were an insufficient number of mitotic cells found in the KD cells to perform an in-depth mitotic defect analysis (**Figure 3F**). The high number of binucleated cells (**Figure 3D**), along with the significantly skewed frequency distribution compared to scramble shRNA KD (**Figure 3F**), suggested cells underwent cytokinetic failure over the 36-hour time period.

**Fig. 3.**
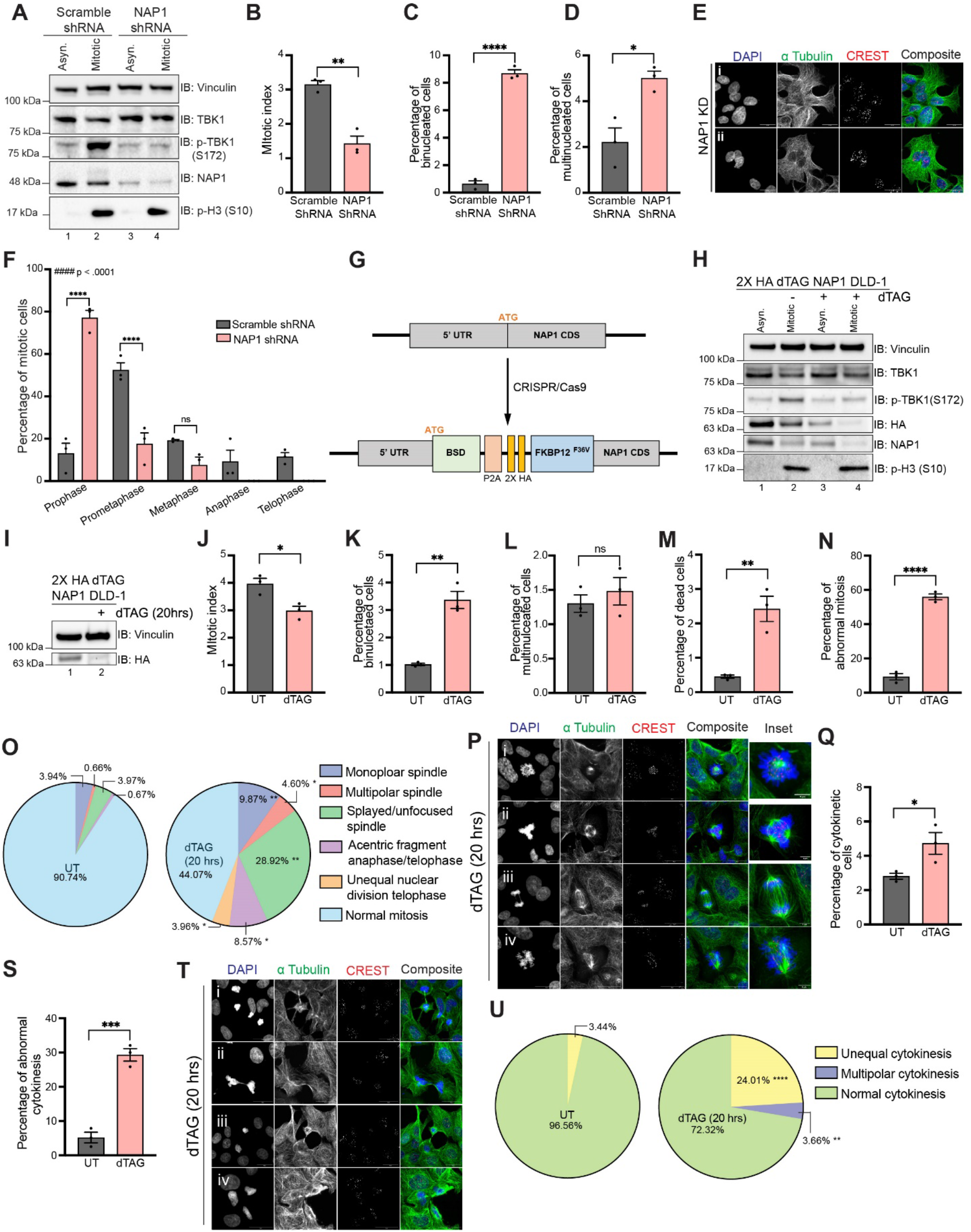
NAP1 loss in a near diploid cell line causes mitotic and cytokinetic defects. **(A**) Western blot analysis of p-TBK1 in asynchronous and mitotic cells from scramble control and transient NAP1 shRNA KD in DLD-1 cells over 36 hours. RO-3306 was used for cell synchronization prior to mitotic release. (**B-D)** Mitotic (B), multinucleated (C), and binucleated (D) cell count percentages from an asynchronous population of scramble control and transient NAP1 KD DLD-1 cells. Error bars indicate ±SEM; n=3 independent experiments. Random fields of view were captured sampling approximately 800 cells per biological replicate from each category. **(E)** Representative confocal images of NAP1 KD cells: (i) binucleated, (ii) multinucleated examples. DAPI (blue) was used as a nuclear counterstain, α-tubulin for cytoskeleton staining (green), and CREST for kinetochore staining (red). Scale bar, 20 μm. **(F)** Mitotic stage frequency distribution of scramble control and NAP1 KD DLD-1 cells. Error bars indicate ±SEM; n=3 independent experiments. Random fields of view were captured sampling approximately 800 cells per biological replicate from each category. Unpaired Student’s t-test was performed for all statistical analysis (B-D, F). For (F), Student’s t-test compared the difference between groups in split in each phase of mitosis. * p < .05, ** p < .01, **** p <.0001, ns = not significant. Kolmogorov-Smirnov nonparametric test was used to analyze the differences between the distribution between groups #### p < .0001. **(G)** Cartoon diagram of dTAG knock-in construct designed to add FKBP12^F36V^ to the N-terminus of NAP1. **(H)** Western blot analysis of p-TBK1 and NAP1 levels in asynchronous and mitotic cells from FKBP12^F36V^-NAP1 DLD-1 cells. RO-3306 was used for cell synchronization prior to mitotic release. Cells were treated with dTAG^V^-1 for 3 hrs in asynchronous conditions and 3 hrs prior and during release in mitotic conditions. **(I)** Western blot analysis of NAP1 levels in asynchronous cells with or without dTAG^V^-1 treatment for 20 hours. (**J-N)** Mitotic (D), multinucleated (E), binucleated (F), dead (G) and abnormal mitotic (H) cell count percentages from an asynchronous population of FKBP12^F36V^-NAP1 DLD-1 cells untreated or treated with 20 hrs of dTAG^V^-1. Error bars indicate ±SEM; n=3 independent experiments. For mitotic index, multinucleated, and binucleated cell counts, random fields of view were captured sampling approximately 800-900 cells per biological replicate from each genotype. For abnormal mitotic cell counts, random fields of view were captured to sample approximately 50 mitotic cells per biological replicate from each genotype. **(O)** Pie charts representing the percentage of different types of mitotic defects found in untreated and dTAG^V^-1 treated (20 hrs) FKBP12^F36V^-NAP1 DLD-1 cells. At least 50 mitotic cells per biological replicate from each genotype were analyzed. n=3 independent experiments. **(P)** Representative confocal images of mitotic defects seen in DLD-1 cells after 20hrs of dTAG^V^-1 treatment: (i) monopolar spindle, (ii) multipolar spindle, (iii) acentric fragment during anaphase, (iv) unfocused spindle poles. DAPI (blue) was used as a nuclear counterstain, α-tubulin for cytoskeleton staining (green), and CREST for kinetochore staining (red). Scale bar, 20 μm, insets, 5μm. **(Q-S)** Cytokinetic (K) and abnormal cytokinetic (L) cell count percentages from untreated and treated (20 hrs) FKBP12^F36V^-NAP1 cells. Error bars indicate ±SEM; n=3 independent experiments. Random fields of view were captured sampling approximately 800-900 cells per biological replicate from each genotype. **(T)** Representative confocal images of cytokinetic defects seen in FKBP12^F36V^-NAP1 DLD-1 cells after 20hrs of dTAG^V^-1 treatment: (i-ii) unfinished and incomplete cytokinesis, (iii) unequal cytokinesis, (iv) multipolar cytokinesis. DAPI (blue) was used as a nuclear counterstain, α-tubulin for cytoskeleton staining (green), and CREST for kinetochore staining (red). Scale bar, 20 μm. Student’s t-test was performed for all statistical analysis. * p < .05, ** p < .01, *** p <.001, **** p <.0001, ns = not significant **(U)** Pie chart representing the percentage of different types of cytokinetic defects found in untreated and dTAG^V^-1 treated (20 hrs) FKBP12^F36V^-NAP1 DLD-1 cells. Random fields of view were captured sampling approximately 30-40 cytokinetic cells per biological replicate from each genotype.

Considering that the KD DLD-1 cells had NAP1 disrupted over a span of two cell divisions, we utilized the degradation tag system (dTAG) [47, 48] for target-specific protein degradation to generate a FKBP^F36V^-NAP1 DLD-1 cell line to allow for immediate and selective manipulation of NAP1 instead of relying on the temporal time scale required for KD efficiency to occur with viral transduction. This would allow us to characterize mitotic defects caused due to NAP1 loss within one mitotic division period and minimize cell death. The FKBP^F36V^ variant allows for selective recognition by a dTAG ligand, like dTAG-13 or dTAG^V^-1, to induce dimerization of the FKBP^F36V^ fused NAP1 to the CRBN or VHL E3 ligase for ubiquitin proteosome degradation (UPS) [47, 48] (**Figure 3G**). After the addition of dTAG^V^-1, NAP1 degradation occurred in both asynchronous and synchronized mitotic cells, and p-TBK1 activation was dampened as compared to untreated mitotic lysates (**Figure 3H**).

To observe the consequences of NAP1 loss in DLD-1 cells over one cell division period in an asynchronous population, we treated the dTAG^V^-1 to FKBP^F36V^-NAP1 cell line for 20 hours (**Figure 3I**). In line with our results when NAP1 was knocked down for 36 hours, we observed a decline in mitotic index and an accumulation of binucleated cells, but multinucleated cell accumulation did not differ (**Figure 3J-L**). The lack of multinucleated cells was not completely unexpected as only one round of cell division occurred. The stress of the sudden degradation of NAP1 did lead to an increase in the number of dead cells present after dTAG^V^-1 treatment (**Figure 3M**). dTAG^V^-1 treated FKBP^F36V^-NAP1 cells had a significantly higher percentage of abnormal mitotic cells present (**Figure 3N**) with many more mitotic defects (**Figure 3O-P**) than seen in the NAP1 KO HeLa cell line (**Figure 2J-K**). This result extended again into cytokinesis where dTAG ^V^-1 treated FKBP^F36V^-NAP1 cells had a higher percentage of cytokinetic cells that were abnormal with significant unequal and multipolar cytokinetic cells (**Figure 3Q-U**). These data suggest NAP1 is a required for both mitosis and cytokinesis most likely through the activation of TBK1.

### TBK1 selectively interacts with and binds NAP1 during mitosis

RNA-seq and northern blotting data demonstrated that NAP1 is expressed ubiquitously in multiple different tissue types [26, 49]. To check the protein level expression in these tissues, we probed a human tissue blot for NAP1, and found it to be expressed in all human tissues except for small intestine, in which there was little detectable NAP1 protein (**Figure 4A**). Vinculin levels were also low for small intestine, but NAP1 protein levels corroborating the previously reported mRNA levels in this tissue [26]. These data confirm NAP1 is a ubiquitously expressed protein most likely important for a wide range of tissue and cell types, supporting the evidence it has a critical function like in cell division.

**Fig. 4.**
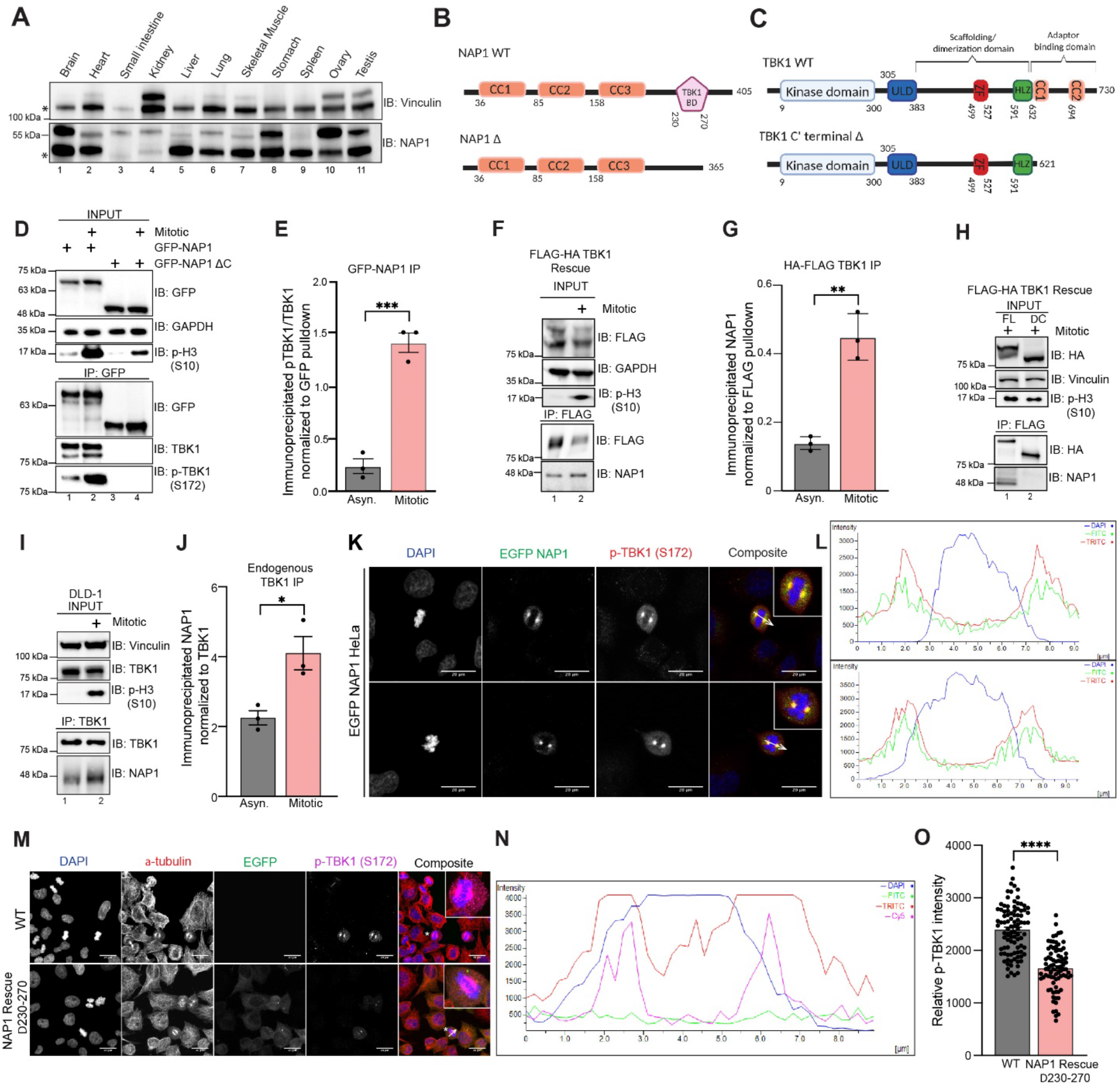
NAP1 binds to TBK1 during mitosis and is localized to centrosomes. **(A)** Western blot analysis of multiple different human tissue samples probed for NAP1. Vinculin was used as a loading control. * Indicates non-specific bands. **(B)** Cartoon representing protein domains of full length (FL) NAP1, and NAP1 lacking the TBK1 binding domain (Δ230-270). **(C)** Cartoon representing protein domains of full length (FL) TBK1, TBK1 lacking adaptor binding domain (Δ C’ terminal). **(D)** Representative immunoblots of the pulldown of transiently expressed GFP-NAP1 or GFP-NAP1 Δ230-270 (lacking TBK1 binding domain) in asynchronous and synchronized mitotic HEK293T cells. Nocodazole was used for cell synchronization. **(E)** Quantification of the ratio of pTBK1/TBK1 signal after normalization to the pulldown efficiency. Error bars indicate ±SEM; n = 3 independent experiments. **(F)** Representative immunoblots of the pulldown of HA-FLAG TBK1 in asynchronous and synchronized mitotic TBK1 rescue HeLa cells. Nocodazole was used for cell synchronization. **(G)** Quantification of the ratio of NAP1 signal after normalization to the pulldown efficiency of TBK1. Error bars indicate ± SEM; n = 3 independent experiments. **(H)** Representative immunoblots of the pulldown of HA-FLAG TBK1 and HA-FLAG TBK1 Δ C’ (lacking adaptor binding domain) in synchronized mitotic TBK1 rescue HeLa cells. Nocodazole was used for cell synchronization. **(I)** Representative immunoblots of the pulldown of endogenous TBK1 in asynchronous and synchronized mitotic DLD-1 cells. Nocodazole was used for cell synchronization. **(J)** Quantification of the ratio of endogenous NAP1 signal after normalization to the pulldown efficiency of TBK1. Nocodazole was used for cell synchronization. Error bars indicate ± SEM; n = 3 independent experiments. **(K)** Representative confocal images of HeLa cells transiently expressing EGFP-NAP1 with immunocytochemical detection of p-TBK1 (red). DAPI (blue) was used as a nuclear counterstain. White arrow depicts area used for line scan analysis in (H). Scale bar = 20μm. **(L)** Line scan analysis of images in (K) generated from Nikon Elements software. **(M)** Representative confocal images of WT HeLa and stable EGFP NAP1 Δ230-270 rescue cells with immunocytochemical detection of p-TBK1 (magenta). DAPI (blue) was used as a nuclear counterstain, and α-tubulin for cytoskeleton staining (red). Scale bar = 20 μm. **(N)** Line scan analysis of images in (M) generated from Nikon Elements software. **(O)** Relative intensity of pTBK1 on centrosomes of mitotic cells from WT HeLa and stable EGFP NAP1 Δ230-270 rescue cells. 40-50 mitotic cells per group were quantified from 2 biological replicates. Error bars indicate ±SEM. Unpaired Student’s t-test was performed for all statistical analysis. * p < .05, ** p < .01, *** p <.001, **** p <.0001

Next, we wanted to determine if there was an interaction between NAP1 and TBK1 during mitosis, as the literature suggests that adaptor binding must occur for TBK1 activation [37, 38]. We performed co-immunoprecipitation (co-IP) experiments by transiently overexpressing either full length GFP-NAP1 (N’ EGFP FL NAP1) or GFP-NAP1 lacking the TBK1 binding domain (N’ EGFP NAP1 Δ230-270) in HEK293T cells [28](**Figure 4B**). Next, we performed the reciprocal co-IP by stably expressing two different N’FLAG-HA-TBK1 rescue constructs (**Figure 4C**) in TBK1 KO HeLa cells as our previous data indicated that TBK1 levels are tightly regulated inside the cell [12]. Phosphorylated endogenous TBK1 was enriched upon the immunoprecipitation of N’ EGFP FL NAP1 with significantly increased binding during mitosis (**Figure 4D-E**). Binding of p-TBK1 was abolished in both asynchronous and mitotic conditions when NAP1 lacked its TBK1 binding domain (**Figure 4D**). Endogenous NAP1 was enriched upon immunoprecipitation of full length TBK1 (N’FLAG-HA FL TBK1) during mitosis (**Figure 4F-G**). This binding did not occur in rescue lines when the known C-terminal TBK1 adaptor binding site was deleted (N’FLAG-HA TBK1ΔC’) (**Figure 4C, H**). Additionally, we verified the enriched binding between NAP1 and TBK1 during mitosis by performing immunoprecipitation of endogenous TBK1 in DLD-1 cells (**Figure 4I-J**).

Next, we sought to corroborate our IP data with experiments to detect the subcellular location of NAP1 during mitosis. Due to the unavailability of antibodies suitable to detect endogenous NAP1 by immunofluorescence, we transiently expressed N’EGFP NAP1 in HeLa cells. We found that NAP1 colocalized with p-TBK1 on the centrosomes of mitotic cells (**Figure 4K-L**). To verify if NAP1 localization on the centrosomes is dependent on TBK1 binding, we stably rescued NAP1 Δ230-270 lacking its TBK1 binding domain in NAP1 KO cells. We failed to observe GFP-NAP1 Δ230-270 localization on the centrosomes during mitosis, and the intensity of p-TBK1 signal in these cells were significantly decreased (**Figure 4M-O**). These data suggest that activation of TBK1 during mitosis is dependent on NAP1 binding.

Although our data suggests that NAP1 has a separate function in regulating TBK1 during mitosis, previous studies of NAP1 have been limited to understanding its role during innate immunity [20, 26, 28, 34]. It is possible that the binding of NAP1 to TBK1 during cell cycle could aberrantly activate innate immune pathways. NAP1 is an adaptor for TBK1 during the innate immune response in which TBK1 on the endoplasmic reticulum phosphorylates the transcription factors, interferon regulator factor 3 (IRF3), causing their translocation to the nucleus resulting in the stimulation of the Type I interferon response [20, 28, 34, 50, 51].

Using a human monocytic cell line that is highly responsive to immunogenic stimuli, we first confirmed that this cell line displayed TBK1 activation during mitosis (**Figure S2A**) and after exposure to innate immune responsive stimuli (LPS and poly I:C) (**Figure S2B**). We then assessed expression of a few select cytokine genes, *Il10, Il6, TNFA*, whose expression is known to be upregulated upon LPS and poly I:C treatment (**Figure S2C-D**). During mitosis, the expression of these genes was either downregulated or only slightly upregulated, in the case of TNF-α, but not reaching the upregulation levels seen during immune stimuli (**Figure S2C-E**) indicating that TBK1 activation during mitosis does not triggering a robust innate immune response. Phosphorylation of either IRF3 or an alternative innate immune kinase (IKKε) that works in conjugation with TBK1 and shares high sequence homology [52], occurred only in response to immunogenic stimuli (LPS and poly I:C) and not during mitosis (**Figure S2F**). Together, these data suggest there exists a distinct mitotic NAP1-TBK1 signaling axis at centrosomes.

### NAP1 protein expression is cell cycle regulated

We continually observed a reduction in NAP1 protein during mitosis across four different cells lines (HeLa, DLD-1, THP1, and RPE-1, an additional near-diploid cell line) independent of the method of synchronization (**Figure 1F, 2A, 2D, 3A, 3H, 5A-B**). This reduction in NAP1 during mitosis also occurred at the mRNA level (**Figure 5C-D**). However, this was not surprising as global transcription is repressed during mitosis [53, 54]. Inhibition of protein synthesis with cycloheximide over 6 hours did not disrupt NAP1 protein levels in asynchronous conditions (**Figure 5E-G**). Also, the inhibition of lysosomal fusion with chloroquine, which blocks autophagy, showed no change to NAP1 levels in asynchronous conditions (**Figure 5H-J**). From these data we conclude that NAP1 is a relatively stable protein except during mitosis.

**Fig. 5.**
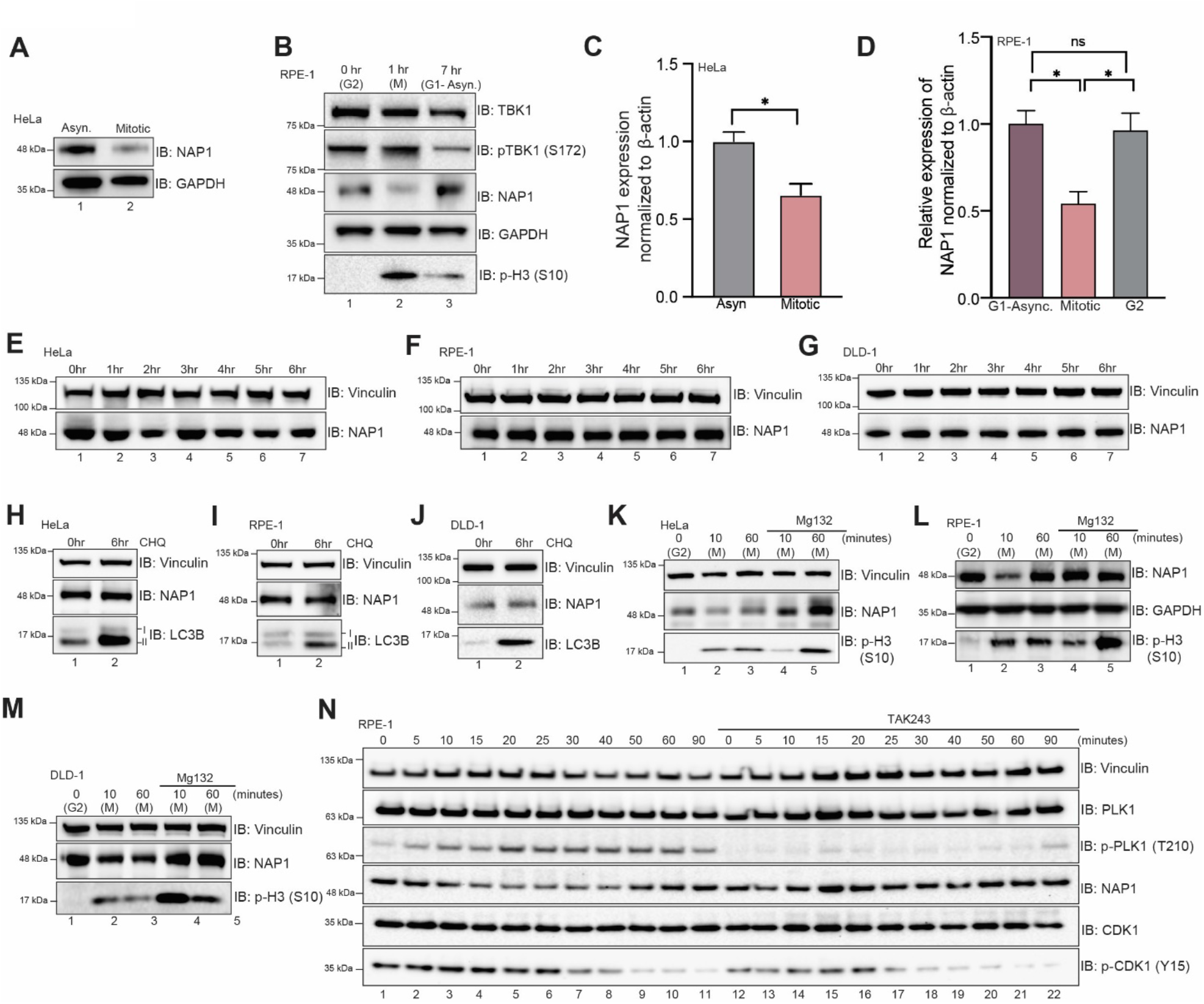
NAP1 protein expression is cell cycle regulated. **(A)** Western blot analysis of NAP1 in asynchronous and synchronized mitotic cells from WT HeLa cells. Nocodazole was used for cell synchronization at mitosis. **(B)** Western blot analysis of NAP1 in G2, mitotic, and G1-asynchrnous RPE-1 cells. RO-3306 was used for cell synchronization prior to mitotic release. **(C)** NAP1 mRNA expression normalized relative to actin in asynchronous and synchronized mitotic HeLa cells. Nocodazole was used for cell synchronization at mitosis. **(D)** NAP1 mRNA expression normalized relative to actin in G2, mitotic, and G1-asyn RPE-1 cells. Error bars indicate ±SD; n = 3 independent experiments. Cells were synchronized at G2 using RO-3306. G2 cells were released for approximately 45-60 minutes to collect mitotic, and for approximately 7 hours to collect G1 samples. (**E-G)** Western blot analysis of NAP1 protein levels after cycloheximide treatment up to 6 hours in HeLa (D), RPE-1 (E), and DLD-1 (F) cells. **(H-J)** Western blot analysis of NAP1 protein levels after 6 hrs of chloroquine treatment in HeLa (G), RPE-1 (H), and (I) cells. (**K-M)** Western blot analysis of NAP1 protein level after MG132 treatment followed by G2 release in HeLa (B), RPE-1 (C), and DLD-1 (D) cells. RO-3306 was used for cell synchronization prior to mitotic release. **(E)** Western blot analysis of NAP1 after TAK243 treatment after G2 release in RPE-1 cells. RO-3306 was used for cell synchronization prior to mitotic release. Student’s t-test or one way ANOVA was performed for all statistical analysis. * p < .05, ns = not significant.

To better define when and how NAP1 levels decreased during mitosis, we synchronized cells using the CDK1 inhibitor RO-3306 at G2 and released cells into mitosis. NAP1 was significantly reduced 10 minutes after release across cell lines (HeLa, RPE-1, and DLD-1), but the protein level mostly recovered during cytokinesis 60 minutes after release (**Figure 5K-M, lanes 1-3**). However, NAP1 levels did not degrade 10 minutes after release from G2 with the addition of the proteasome inhibitor, MG132 (**Figure 5K-M, lanes 4, 5)**. This indicates that the UPS was involved in this reduction of NAP1 during mitosis. To better characterize the exact time-point at which NAP1 levels degrade during mitosis and return to levels seen during asynchronous-G1 conditions, we used the RPE-1 cell line. Using p-PLK1 T210 and p-CDK1 Y15 as markers to indicate different stages during mitosis, we saw that NAP1 levels continuously degrade up until 40 minutes (anaphase) after G2 release (**Figure 5N, lane 8**). Levels began to increase and recover at the 50- and 60-minute time points (cytokinesis), and 90-minute time point (asynchronous, G1) (**Figure 5N, lanes 9-11**). We treated another set of cells with the E3 activating enzyme inhibitor TAK243 as an alternative method to inhibit the UPS [55] and saw that NAP1 levels were stabilized throughout the time points collected after G2 release (**Figure 5N**). These data indicate that levels of NAP1 are tightly regulated during mitosis by the UPS.

### Quantitative phospho-proteomics pipeline identifies TBK1 downstream substrates upon mitosis

The complete landscape of proteins targeted by TBK1 during mitosis is unidentified, and the substrates that have been identified do not fully explain the diverse array of mitotic and cytokinetic defects displayed due to the loss of TBK1 or NAP1 (**Figure 2, 3, S1**), nor do they provide insight into the regulation of these proteins [11, 17, 33]. Therefore, using unbiased quantitative phospho-proteomics, we sought to identify additional targets of TBK1 during mitosis. We synchronized WT HeLa and two independently created TBK1 KO HeLa cell lines from our previous work[12] into mitosis. Additionally, to account for off-target gene-editing effects, we also blocked TBK1 pharmacologically with a specific inhibitor, MRT67307 [56] (**Figure 6A**). Thus, phosphopeptides of interest should not be regulated with the small molecule inhibitor in either condition lacking functional TBK1 activity. While some previous studies [5, 6] have quantitatively detailed the global cell cycle and mitotic phosphorylation landscape, to our knowledge, an in-depth, focused quantitative analysis of TBK1-dependent changes in the phosphoproteome during mitosis has not been performed.

**Fig. 6.**
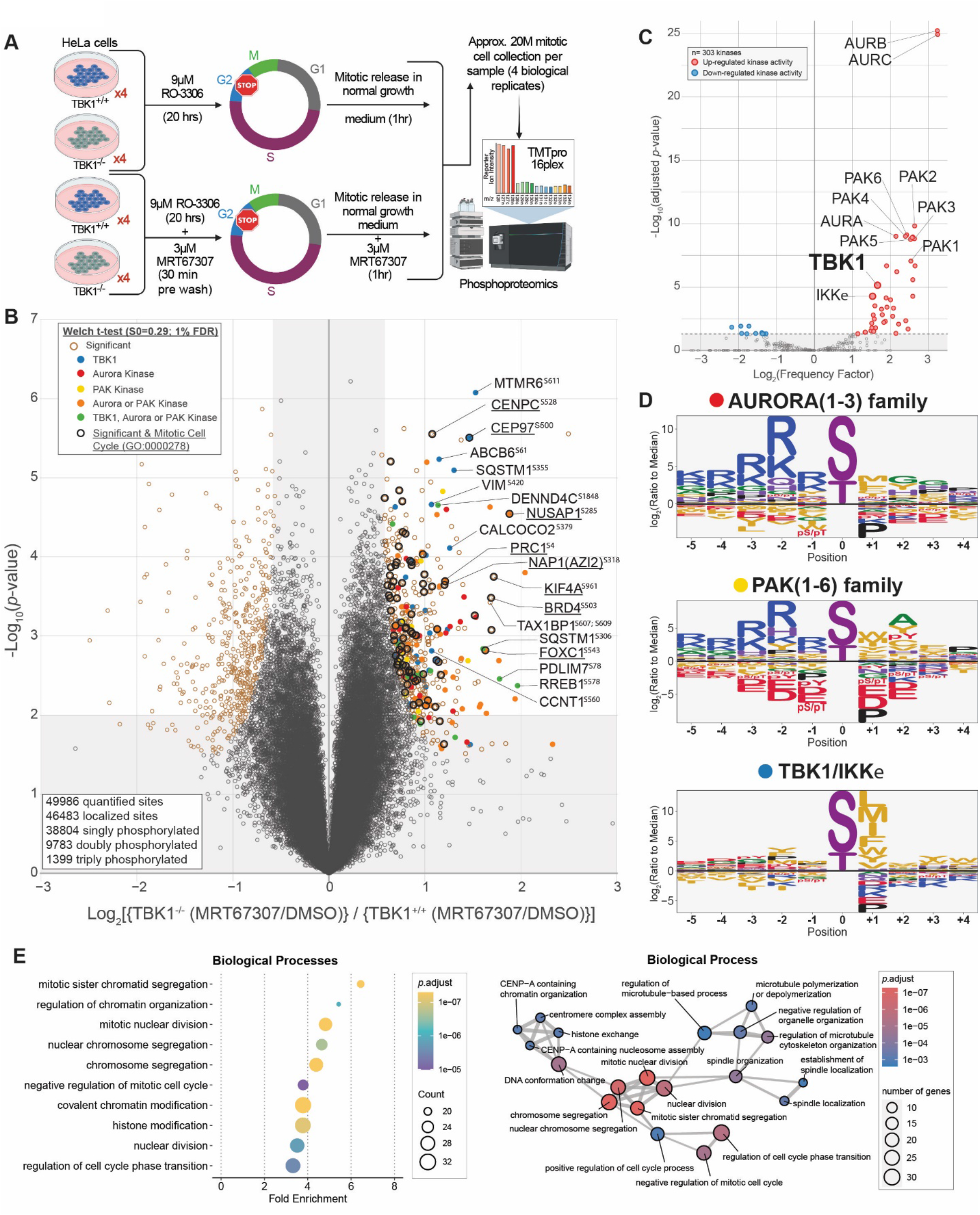
Quantitative phospho-proteomics pipeline identifies TBK1 downstream substrates upon mitosis. **(A)** Workflow for TMTpro-based phosphoproteomics of approximately 20M (million) mitotic cells. 16plex proteomics was performed in quadruplicate on samples harvested 1-hour post-mitotic release. **(B)** Volcano plots [Log_10_ (*p*-value) versus Log_2_ ratio] for representing phosphorylation sites that are affected by MRT67307 and loss of TBK1. Proteins are shown in black circles (non-significant) and red circle (Tier 1 significant). Circles were color coded for the motif that most likely fit TBK1 (blue), Aurora kinases (red), PAK kinases (yellow), Aurora or PAK kinases (orange), or all three (green). Bolded black circles categorize known mitotic proteins (GO:0000278). The inset indicated additional color coding for the statistical analysis. **(C)** Volcano plot scoring each identified substrate against the 303 kinase motifs panel from (Johnson et al., 2022) to determine statistically significant enriched activated kinases. **(D)** Motif analysis using the pLogo tool (plogo.uconn.edu) identifies the motif for the substrate candidates based on synthetic peptide analysis. The y-axis is the log odds of the binomial probability. **(E)** Gene Ontology terms enrichment analysis among the Biological Processes for enriched phosphosites Tier 2. Mitotic and cell cycle related classes were significantly enriched (left panel). Enrichment map networks, each node represents a gene set (i.e., a GO term) and each edge represents the overlap between two gene sets (right panel).

To broadly understand how the phosphoproteome is remodeled and to search for potential mitotic TBK1 substrates, we performed quantitative proteomics on synchronized and released total cell extracts using 16plex TMTpro (**Figure 6A**). Tryptic peptides from whole-cell extracts were subjected to phosphopeptide enrichment, and samples were analyzed using 16plex TMTpro workflow [57], with phosphopeptide intensities normalized with total protein abundance measured in parallel. In total, we quantified 8,650 proteins and 49,986 phosphopeptides in 7150 phosphoproteins from the phosphopeptide enrichment (**Data file S1, S2**). Total proteomics carried out in parallel to normalize phosphopeptides reveal no major stabilization or degradation of protein upon TBK1 loss of function (**Figure S3A**). Principal component analysis revealed reproducible replicate data, with 39.7% of the variance being driven by cell genotype and 27% driven by the small molecule inhibitor treatment (**Figure S3B**). Given that both CRISPR editing and small molecule inhibitors can independently result in off-target effects, we set to compare the four different experimental conditions such that only variations that are abolished in both the CRISPR edited cells and cells treated with MRT67307 would be enriched in our TBK1-focused analysis. From the ∼50,000 unique phosphorylation sites quantified; we identified 63 sites in 56 proteins whose abundance was statistically increased by >2-fold (p < 0.01, 1%FDR) (**Figure 6B**). Several significant sites are previously reported bona fide substrates of TBK1 (e.g., SQSTM1, TAX1BP1), thus validating our experimental approach. Further analysis revealed several enriched sites linked to the mitotic cell cycle gene ontology term, which led us to consider them strong TBK1 substrate candidates.

Considering that the disruption of one kinase can have many subsequential changes to other kinases, phosphoproteomic data although informative, cannot always be traced back to one particular kinase. In this case, we compared our phospho-proteomic data and the motifs surrounding our identified substrates to data generated from the Cantley laboratory that screened all 303 kinases with a synthetic peptide library to determine their motifs [58]. This would enable us to identify if other kinases were potentially affected by the loss of TBK1 as well as sort out which potential substrates were directly TBK1 dependent. Performing a motif analysis using this methodology to determine any enrichment at the sequence level for sites statistically increased by >2-fold (p<0.05, 5%FDR) found that Aurora, PAK, and TBK1/IKKe kinase motifs were predominately represented from our substrate list (**Figure 6B-D, S3C, S4A**).

To further evaluate the global effect of impairing TBK1’s activity upon mitosis, we performed functional enrichment analysis based on Gene Ontology (GO) annotations for Biological Processes (**Figure 6E, S4B**), Molecular Function (**Figure S4C**), and Cellular Component terms (**Figure S4D**). This analysis revealed strong enrichment of many biological processes linked to mitotic cell cycle and chromosome segregation (**Figure 6E, left panel**) and visualized as enrichment map networks (**Figure 6E, right panel**). Taken together, these data assert a role for TBK1 activity in mitotic progression, and we highlight a list of phosphorylation sites and possible TBK1 substrates or indirect substrates (**Figure 6B, Data file S3, S4A**).

We then experimentally validated if Aurora A and B kinases were regulated by TBK1. In DLD-1 and RPE-1 cells treated with TBK1 pharmacological inhibitors, the trans-autophosphorylation sites (T288 AurA, T232 AurB) indicative of activity for Aurora A and B were abolished as well as the downstream p-HistoneH3 S10 site, which is directly downstream of Aurora B [59] (**Figure S4E-F**). TBK1 KO HeLa cells displayed a less prominent reduction, but still displayed a decrease in p-AurA T288 and p-AurB T232 (**Figure S4G**). To our knowledge, this is the first time TBK1 has been implicated in regulation of Aurora A and B. In addition to the effects that TBK1 can have on direct substrates during mitosis, this dysregulation of Aurora kinases would cause defects not only in mitosis but also during cytokinesis, which may better explain the diverse cell division defects we found.

Our GO analyses indicated that microtubule polymerization and function were highly represented. Performing immunofluorescence staining for p-TBK1, the signal colocalized to centrin and γ-tubulin foci during mitosis across two independent cell lines (**Figure S5A-B**) reconfirming that TBK1 is activated on centrosomes. However, the p-TBK1 signal also appeared to be localized to the spindle assembly during metaphase (**Figure S5A-B**). When we compared the coverage area of the p-TBK1 signal, the area is significantly reduced upon the loss of NAP1 or in the TBK1 binding deficient NAP1 rescue line compared to WT cells (**Figure S5C-D**). These results indicate that TBK1 is involved in both centrosomal organization and function, and microtubule processes as indicated by the substrates found in our phosphoproteomic screen.

### TBK1 phosphorylation of NAP1 at S318 impacts its stability during mitosis

Our phosphoproteomics data indicated that NAP1 is phosphorylated at serine 318 during mitosis by TBK1 (**Figure 6B**). Although, the NAP1 sequence motif at this serine residue did not perfectly match the preferred *in vitro* TBK1 sequence motif (**Figure 6D, S3C**), our data suggested it was a bona fide substrate. First, TBK1 and NAP1 are bound, colocalizing at the centrosomes during mitosis (**Figure 4**). Second, TBK1 phosphorylates its own adaptors to modulate their function in other contexts [21, 32, 43]. Third, when comparing the NAP1 sequence motif around S318 against all other 303 kinases, NAP1 still had a high 82.43% kinase preference probability (**Data file S4**). Therefore, we decided to further investigate the connection between NAP1 and TBK1.

First, we tested whether TBK1 affected NAP1 levels. We collected DLD-1 cells at different stages of the cell cycle and checked for NAP1 levels with or without MRT67307 treatment (TBK1 inhibitor). As expected, the control cells had a significant reduction of NAP1 during mitosis compared to asynchronous conditions; however, MRT67307 treated cells had significantly increased levels of NAP1 **(Figure 7A-B)**. Next, we sought to validate NAP1 phosphorylation during mitosis by Phos-tag gel analysis. A higher molecular weight band indicating the presence of a phosphorylated form of NAP1 appeared, which was missing from either TBK1 inhibitor treated or phosphatase treated mitotic cell lysates **(Figure 7C)**. NAP1 stability was also significantly increased during mitosis in a TBK1 kinase dead mutant (K38A) rescue line **(Figure 7D-E)**. The mitotic lysates from TBK1 K38A cells displayed a reduced higher molecular weight NAP1 banding pattern indicating TBK1 K38A was not able to phosphorylate NAP1 (**Figure 7F**). We then generated S318A phospho-deficient NAP1 rescue line to test whether TBK1 phosphorylation affected NAP1 stability during mitosis. NAP1 S318A resulted in a significantly higher amount of NAP1 during mitosis as compared to asynchronous conditions (**Figure 7G-H**). We also confirmed this result in our TBK1 KO line where NAP1 appeared less degraded during mitosis (**Figure 7I**). Together, these data suggest that activated TBK1 phosphorylates NAP1 on serine 318, which regulates NAP1 degradation **(Figure 7J)**.

**Fig. 7.**
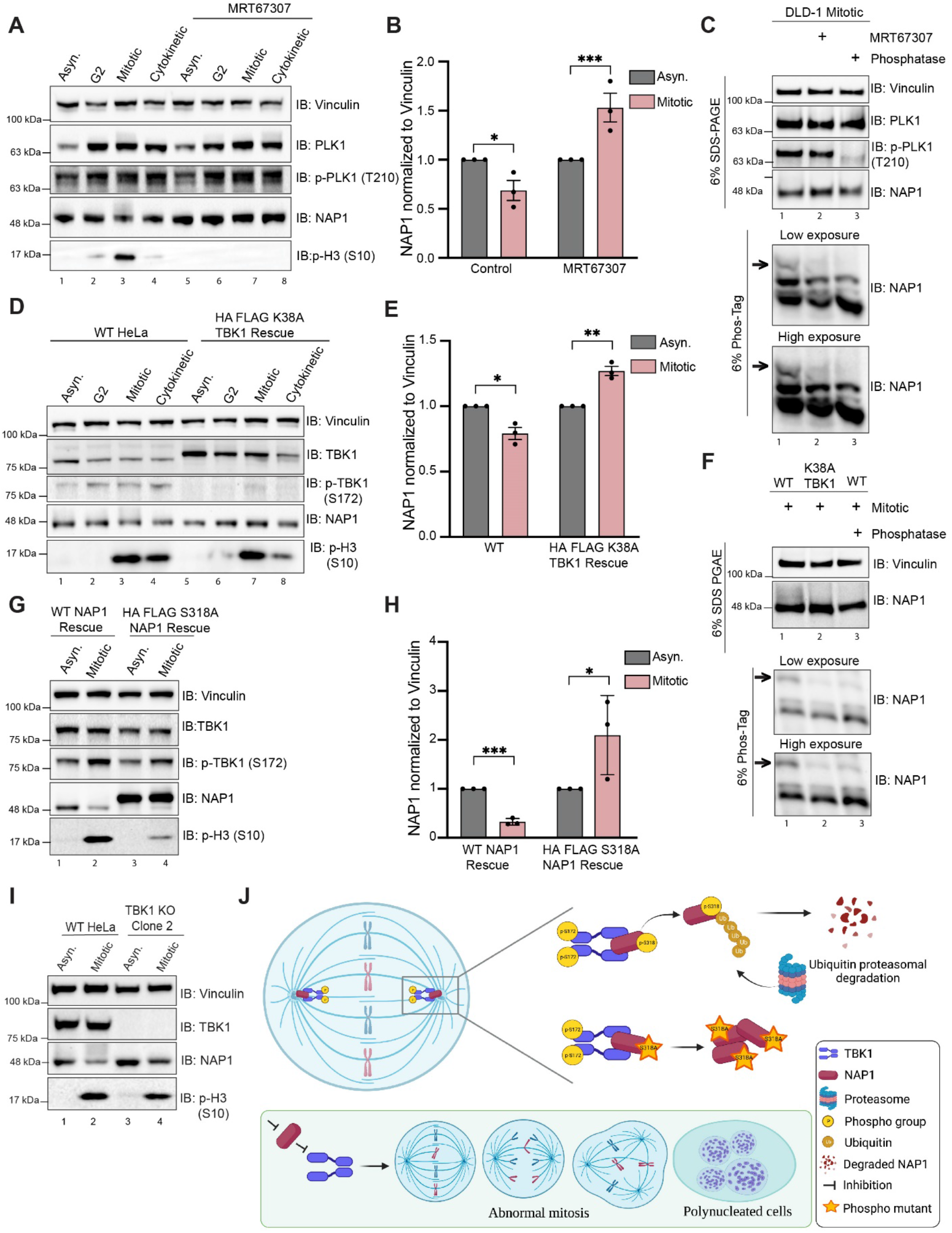
TBK1 phosphorylation of NAP1 at S318 impacts its stability during mitosis. **(A)** Representative immunoblots of asynchronous, G2, mitotic, and cytokinetic DLD-1 cells with or without TBK1 inhibitor treatment (MRT67307) for 1 hr prior to G2 and throughout release timepoints. Asynchronous cells were treated for 2 hrs. Cells were synchronized at G2 using RO-3306. G2 cells were released for approximately 30-45 minutes to collect mitotic samples and for approximately 80-90 minutes to collect cytokinetic cells. **(B)** Quantification of normalized NAP1 protein levels during asynchronous and mitotic conditions with or without MRT67307. Error bars indicate ±SEM; n=3 independent experiments. **(C)** Phos-Tag gel analysis of NAP1 mitotic protein from DLD-1 cells either with MRT67307 for 1 hr prior to G2 and throughout release or with 1 hr phosphatase treatment. 6% SDS-PAGE gel was run in tandem to ensure equal protein loading. **(D)** Representative immunoblots of asynchronous, G2, mitotic, and cytokinetic cells from WT HeLa and stable TBK1 K38A rescue lines. Cells were synchronized at G2 using RO-3306. G2 cells were released for approximately 30-45 minutes to collect mitotic samples and for approximately 80-90 minutes to collect cytokinetic cells. **(E)** Quantification of normalized NAP1 protein levels during asynchronous and mitotic conditions in WT HeLa and stable TBK1 K38A rescue lines. Error bars indicate ±SEM; n=3 independent experiments. **(F)** Phos-Tag gel analysis of NAP1 mitotic protein from HeLa, K38A TBK1 rescue, and HeLa cells treated 1 hr with phosphatase. 6% SDS-PAGE gel was run in tandem to ensure equal protein loading. **(G)** Representatives immunoblot analysis of NAP1 in asynchronous and mitotic cells from stable NAP1 and NAP1 S318A rescue lines. Cells were synchronized using RO-3306 prior to mitotic release. **(H)** Quantification of normalized NAP1 protein levels during asynchronous and mitotic conditions in stable NAP1 rescue, and stable NAP1 S318A rescue lines. Error bars indicate ±SEM; n=3 independent experiments. **(I)** Western blot analysis of NAP1 levels in HeLa and TBK1 KO cells in asynchronous and mitotic cells. Nocodazole was used for cell synchronization at mitosis. **(J)** Cartoon representing mechanistic regulation of NAP1 and TBK1 during mitosis. NAP1 activates TBK1 on centrosomes by binding to its adaptor binding C’ domain. Activated TBK1 phosphorylates NAP1 on S318 which acts as a signal for ubiquitin proteasomal degradation of NAP1. Loss of either of these centrosomal proteins, NAP1 or TBK1, impairs mitosis and cytokinesis leading to retention of multinucleated cells. Unpaired Student’s t-test was performed for all statistical analysis. * p < .05, ** p < .01, *** p <.001.

## Discussion

Our work demonstrates that NAP1/AZI2 is a mitotic and cytokinetic regulatory protein required for normal cell cycle progression. Mechanistically, NAP1 binds to TBK1 at centrosomes to activate TBK1, which subsequently phosphorylates several mitotic and cytokinetic proteins which we identified. NAP1 protein levels during mitosis are tightly regulated by the ubiquitin proteasome system where TBK1 phosphorylation of NAP1 at S318 influences its stability, thus acting as a possible feedback regulatory mechanism (**Figure 7J**).

Previous studies including our own have not extensively characterized all the consequences to cell division upon the loss or pharmacological inhibition of TBK1, although these studies reported an increase in the number of multinucleated cells [11, 12, 17]. Previous experiments treating different cancer cell lines with TBK1 inhibitors or siRNAs showed an increase in the presence of supernumerary centrosomes and misaligned spindle poles [11, 17]. However, our results find that these defects are extended much further than previously thought, affecting every stage of mitosis and cytokinesis. Mitotic and cytokinetic analyses with cells lacking either TBK1 or NAP1 displayed many of the same phenotypes, which we would expect if NAP1 and TBK1 act within the same pathway. Furthermore, we also utilized in this study a near diploid cell line amendable to mitotic studies, which has not previously been performed. In agreement with the previous study utilizing TBK1 pharmacological inhibitors in cancerous transformed cells [11], transient KD of NAP1 in DLD-1 cells over 36 hours could not progress past metaphase hindering an extensive mitotic characterization of these cells.

The plethora of mitotic and cytokinetic defects found in both NAP1 and TBK1 KO cells led us to perform phosphoproteomic analysis to unbiasedly discover additional TBK1 substrates to explain these diverse types of defects. Comparing the phosphorylation sites to the kinome substrate specificity motifs defined by synthetic peptide libraries [58], substrates identified indicated that TBK1 either directly or indirectly affects Aurora A/B and PAK kinase activity as their motifs were highly represented. This was in addition to substrates that are direct TBK1 substrates. In DLD-1 and RPE-1 cells treated with TBK1 pharmacological inhibitors, Aurora A and Aurora B kinase activity was abolished. However, HeLa cells lacking TBK1 did not have such a dramatic reduction in p-Aurora A T288 and p-Aurora B T232 likely due to its state of aneuploidy compensating in some unknown manner. These results also provide an explanation as to why we were unsuccessful in generating a NAP1 KO or stable KD cell line and instead had to generate the FKBP^F36V^-NAP1 knock in line to characterize mitotic defects when cells were near diploid. The choice to utilize different TBK1 KO clones with and without the TBK1 pharmacological inhibitor was to increase stringency in our experimental design to avoid the detection and identification of non-specific phosphorylation sites. However, the generation of an inducible conditional TBK1 knockout diploid cell line may better distinguish phosphor-sites in substrates that are compensated for in other cell lines. In this case, the balance between temporal timing of the loss of the protein balanced with the concern for cell death may need to carefully need to be considered. This also may explain why a previous substrate Plk1 [33], which was identified in an asynchronous TBK1 KD screen in A549 cells, did not appear as a major hit nor did Plk1 activity appear disrupted in our DLD-1 MRT67307 treated cells.

Regardless, to our knowledge, this is the first time TBK1 has been implicated in AurA and B kinase activity and does explain how loss of TBK1 and NAP1 is so detrimental. Previously reported substrates, NuMa, CEP170, Plk1, Cdc20, Cdh1, all function during different stages of mitosis [11, 17, 33], but these substrates do not explain all of the cytokinetic defects observed or the increase in binucleated cells in NAP1 and TBK1 KO cells, which can be indicative of cytokinetic failure. AurB is important in the formation of the contractile ring and cleavage furrow during cytokinesis [60-62], so its disruption alone may explain these defects upon the loss of TBK1 and NAP1. Future work will be required to determine the mechanism by which TBK1 affects these other kinases and the careful dissection between bona fide TBK1 substrates and those of other kinases TBK1 regulates.

The *drosophila* orthologue of TBK1, ik2, is associated with the minus ends of microtubules [63]. *ik2* mutant flies display bristle and oocyte abnormalities due to defects in microtubule and cytoskeletal organization in part by its inability to phosphorylate spn-F and correctly localize ik2 to microtubules [63-67]. Microtubule disruption activates ik2, but dominant negative ik2 reduces microtubule polymerization suggesting its role in increasing microtubule stability [68]. We observed p-TBK1 confined with g-tubulin and centrin during mitosis, but its localization appears to be on the spindle assembly excluding the astral microtubules. TBK1 phosphorylating NuMa by way of its interaction with CEP170 may be responsible for its association to the spindle poles and correct microtubule tethering [11]. However, another paper in the context of Zika infection suggested that CEP63 is involved in TBK1 recruitment to centrosomes identified by g-tubulin and centrin [69]. Considering that there is no mammalian orthologue to *spn-F* and our analysis revealed several microtubule-associated proteins largely represented in our proteomics data, more work is required to determine specifically where TBK1 is active or inactive on microtubules during different cellular contexts and what downstream consequences this may have to microtubule stability.

We attribute the ability for NAP1 to regulate TBK1 due to its ability to bind to the C’ terminal of TBK1. These results are not surprising as it has previously been shown that multiple adaptors bind to this region of TBK1 to facilitate dimerization and trans-autophosphorylation of the S172 site to activate kinase activity [37, 38]. However, surprisingly, our data suggests that NAP1 localization to the centrosomes is dependent on TBK1 binding, which is opposite of previous findings. TBK1 adaptors are thought to facilitate TBK1 translocation and localization to target damaged mitochondria, bacteria, or exogenous innate immune triggering stimuli [20, 31, 32, 42]. In our observations, NAP1 no longer localizes to centrosomes when lacking its TBK1 binding domain (NAP1 D230-270) nor is TBK1 mis localized upon the loss of NAP1. The area and intensity of p-TBK1 signal is weaker but activated TBK1 is still localized at the centrosomes. This indicates that TBK1 may translocate in a manner independent of its adaptors during cell division. Another possibility is that there is another adaptor not yet identified that can also activate TBK1 during mitosis and participate in regulating its localization or an adaptor that normally does not participate in mitosis can compensate to activate TBK1 in response to the loss of NAP1. TBK1 KO mice are embryonic lethal at day 14.5 [16]. However, the NAP1 knockout mouse is viable, but does display proliferative defects in GM-dendritic cells [70]. We observed the presence of other TBK1 adaptors being phosphorylated during mitosis, but their role is unknown. How TBK1 localizes to centrosomes in the first place is quite intriguing and requires future study.

TBK1 expression is tightly regulated, and our previous data showed abnormal activation and subcellular localization upon overexpression of TBK1 [12]. Elevated expression of TBK1 has been reported in many different types of cancer with high expression correlated with negative patient outcomes [13-15] This may explain our data showing the tight regulation of NAP1 expression which displays UPS dependent degradation from prophase to anaphase, but expression begins to increase and normalize to baseline in the late phases of mitosis and cytokinesis. The regulation of NAP1 protein levels during mitosis may ensure that TBK1 is not abnormally activated or overactivated, nor persists longer than necessary during mitosis. To understand how the degradation of NAP1 affects TBK1 activation and function during mitosis, manipulation of the E3 ligase responsible for the ubiquitination and degradation of NAP1 is necessary. Future studies for E3 ligase screens are warranted to identify this protein.

We characterized the role of NAP1 and TBK1 during mitosis; however, other reports indicate that genetic influence and external environmental signaling/stimuli also influence cell division in a TBK1-dependent manner. TBK1’s role in cell proliferation was originally described in KRAS mutant cell lines [71], but KRAS mutations interacting with TBK1 are not the sole driver of the proliferation defects [11]. Proliferation stimulated by growth factor signaling is reliant on SINTBAD for the activation of TBK1 rather than NAP1 [72]. The subcellular localization of TBK1 and its adaptors in these genetic and environmental contexts as well as TBK1 substrates in these situations have yet to be described, making it uncertain if described differences in proliferation are due to changes in mitosis and/or cytokinesis. Both growth factor signaling and KRAS mutations converge on ERK kinase signaling pathways[73, 74], and sustained ERK activation throughout G1 is required for S phase entry [75-77]. Thus, it is possible that TBK1 can affect other stages of the cell cycle.

While NAP1 was first identified as an innate immune adaptor [26], our data supports a NAP1-TBK1 activation axis occurring at centrosomes ascribing it a new role as a regulator of mitosis and cytokinesis. We have provided mechanistic insight into its function during cell division by providing evidence of how NAP1 activates TBK1 and provided data on how its own stability may be negatively regulated in a TBK1-dependent manner. Most research studies on TBK1-dependent processes such as mitophagy and innate immunity concentrate on asynchronous cell populations or post-mitotic cells ignoring the interplay between cell division and these other processes. However, the ubiquitous nature of TBK1 signaling across different cell types indicate these considerations are warranted. Future work to understand how innate immunity, selective autophagy, and cell division intersect is highly attractive as all three are implicated in human diseases such as cancer.

## Materials and Methods Cell culture

HeLa and HEK293T cells were maintained in DMEM high glucose medium supplemented with 10% FBS, 2mM L-Glutamine, 10mM HEPES, 0.1 mM non-essential amino acids, and 1mM sodium pyruvate. RPE-1 and DLD-1 cells were a kind gift from Dr. Daniela Cimini’s lab. RPE-1 cells were maintained in DMEM/F-12 medium supplemented with 10% FBS. DLD-1 cells were maintained in RPMI-1640 medium with 10% FBS. Penta KO and DKO HeLa cells were a kind gift from Dr. Richard J. Youle’s lab. THP1-Lucia™ ISG cells (Invivogen) were maintained in RPMI-1640 medium with 10% FBS, 2 mM L-glutamine, 25 mM HEPES, 100 μg/ml Normocin™, and Pen-Strep (100 U/ml-100 μg/ml). Cells were routinely tested for mycoplasma contamination by PCR (Southern Biotech).

### Antibodies

The following antibodies were used for this study: TBK1/NAK (#3504S/#14590S; Cell Signaling Technology (CST)), pTBK1 (Ser172; #5483S; CST), pTBK1 Alexa Fluor 488 or 647 conjugated (Ser172; #14586/#14590; CST), NAP1/AZI2 (#15042-1-AP; Proteintech™), GAPDH (G9545; Sigma), Vinculin (#700062; Invitrogen), p-Histone H3 (S10; #53348S; CST), SINTBAD (#8605S; CST), TANK (#2141S; CST), FLAG M2 (#F1804; Sigma), p62 (#H00008878-M01; Abnova), NBR1 (#H00004077-M01; Abnova), TAX1BP1 (HPA024432;Sigma), Optineurin (#10837-1-AP; Proteintech), NDP52 (60732; CST), α-Tubulin (#T6074; Sigma)(#2144S; CST)(ab52866;Abcam), GFP (Cat#11814460001; Roche), IKK□□ (#2905S; CST), pIKK11 (S162; #8766S, CST), HA.11 (#901513; BioLegend), IRF3 (#4302; CST), p-IRF3 (S386; ab76493; Abcam), Plk1 (#4513T; CST), p-Plk1 (Thr210; # 5472T; CST), CREST (#15-234; Antibodies Inc.), CDK1/CDC2 (#77055; CST), pCDK1/CDC2 (Tyr15; #4539; CST), LC3 (NB600-1384; Novus)(#2775S; CST), g-tubulin (#5326; Sigma), Aurora A (#14475; CST), Aurora B (#3094T; CST), Aurora A/B (T288/T232; #2914T; CST), Centrin (#04-1624; EMD Millipore).

### Additional chemicals

The following chemicals were used for this study: 10ug/ul cycloheximide (AC357420010; Fisher Scientific), 10μm MG132 (NC9937881; Fisher Scientific), 1 μM TAK243 (30108; Cayman Chemical), 50 μM chloroquine diphosphate (C2301100G; Fisher Scientific), 3μM MRT67307 (inh-mrt; Invivogen), 5mg/mL LPS (eBioscience), 1 mg/mL poly I:C (Fisher Scientific), 5U FastAP Thermosensitive Alkaline Phosphatase (EF0651; ThermoFisher), 1 μM dTAG^V^-1 (Toris), and10μM BI605906 (50-203-0195; Fisher Scientific).

### Plasmids and constructs

To generate NAP1 rescue lines, NAP1 cDNA was cloned into pDONR223 and transferred into the pHAGE-N’-FLAG-HA-IRES-puro, pHAGE-N’-EGFP-Gaw-IRES-Blast, or pHAGE-C’-EGFP-Gaw-IRES-Blast vectors using LR recombinase (Invitrogen). The pHAGE-N’-FLAG-HA-TBK1 cloning has been described previously [12] and deposited to Addgene #131791. The pHAGE-N’-FLAG-HA-TBK1 K38A construct was a kind gift from Dr. Richard Youle’s lab. The following site directed mutagenesis primers were used to generated mutant constructions: NAP1 Δ230-270: TTC ATC AAG TGC AGT TTT GTA TAT GGA TCC GTT TGT TTG GCT TTC, GAA AGC CAA ACA AAC GGA TCC ATA TAC AAA ACT GCA CTT GAT GAA; TBK1 Δ C’ terminus: GGG GAC AAG TTT GTA CAA AAA AGCAGG CTT CG AGG AGA TAG AAC CAT GAT GCA GAG CAC TTC TAA TCA TCT G, GGG GAC CAC TTT GTA CAA GAA AGCTGG GTC CTA CTA TAT CCA TTC TTCTGA CTT ATT. NAP1 S318A: AATCCTCCAAGCATGGACAGACA, GCTTTCTCTGATAAAACCTTTACATC. All constructs were confirmed by DNA sequencing and deposited on Addgene.

### Retrovirus and lentivirus generation

Dishes were coated with 50μg/mL poly-d-lysine (Sigma); and HEK293T cells were plated at 70-80% confluency before transfection. Lentiviral helpers and constructs were transfected using XtremeGENE 9™ (Roche) according to the manufacturer’s instructions at a 1:3 ratio. 24 hrs after transfection, media was changed. Infectious media containing virus was collected 40 hrs later and filtered with a 0.45μm PES membrane filter (Millipore). Viral purification was performed using an Optima MAX-XP ultracentrifuge (Beckman) and spinning media at 100,000 x *g* for 2hr at 4°C. Viral pellet was resuspended in sterile PBS, and tier was quantified using qPCR Lentivirus Titer Kit (abm) according to the manufacturer’s directions. Live filtered virus was used to transduce cells with polybrene (10μg/ml, Sigma).

### CRISPR knockout cell line generation

In brief, CRISPR design was aided by publicly available software provided by MIT at www.crispr.mit.edu. CRISPR oligos for NAP1: AAA CCA GCT GGA GGA GTT CTA CTT C, CAC CGA AGT AGA ACT CCT CCA GCT G. Primers were annealed with Phusion DNA polymerase (Thermo Fisher Scientific) using the following conditions: 98°C for 1’, 2-3 cycles of (98°C for 10”, 53°C for 20”, 72°C for 30”), 72°C for 5.’ The annealed primers were cloned into the linearized gRNA vector gRNA, which was a gift from Dr. Feng Zhang (Addgene plasmid #62988) using the Gibson Assembly Cloning Kit (NEB). HeLa cells were cotransfected using XtremeGENE 9™ (Roche) using the above CRISPR plasmid. Cells were selected by puromycin (1mg/ml) and serially diluted into 96 well plates to select for single colony clones. DNA was extracted from individual clones using the Zymo gDNA Isolation Kit and genotyped/sequenced using the following primers: Exon 4 F GAAGCGAATGACATCTGCA, Exon 4 R CCTCTTCTGCTTCATCACAACCT.

### shRNA cell line generation

pLKO.1 puro was a gift from Dr. Bob Weinberg (MIT) (Addgene plasmid # 8453) digested with AgeI and EcoRI for 4 hrs at 37°C. Digested plasmid was excised, and gel purified with GeneJET™ gel extraction kit (Thermo Fisher). Oligos were designed for the following target sequences and annealed (NAP1: CCG GCC ACT GCA TTA CTT GGA TCA ACT CGA GTT GAT CCA AGT AAT GCA GTG GTT TTT G; AAT TCA AAA ACC ACT GCA TTA CTT GGA TCA ACT CGA GTT GAT CCA AGT AAT GCA GTG G, TANK: CCG GCC TCA AAG TCT ACG AGA TCA ACT CGA GTT GAT CTC GTA GAC TTT GAG GTT TTT G; AAT TCA AAA ACC TCA AAG TCT ACG AGA TCA ACT CGA GTT GAT CTC GTA GAC TTT GAG G, SINTBAD: CCG GCC TCT GCC TTT CTG TTC TTA ACT CGA GTT AAG AAC AGA AAG GCA GAG GTT TTT G; AAT TCA AAA ACC TCT GCC TTT CTG TTC TTA ACT CGA GTT AAG AAC AGA AAG GCA GAG G). Annealed oligos were ligated into the digested vector with T4 ligase (NEB); and colonies were screened by sequencing. Scramble shRNA was a gift from Dr. David Sabatini (MIT) (Addgene plasmid # 1864). Cells were selected for using 1mg/mL puromycin.

### FKBP12^F36V^ degradation tag-NAP1 cell line generation

The cell line was generated as previously described [78]. For donor vector, the BSD-P2A-2xHA-FKBP12^F36V^ and backbone pCRIS-PITCH cassettes were generated using PCR. The NAP1 homology sequences for N terminal tagging were synthesized as a gBlock gene fragment (IDT) and they were assembled using NEBuilder HiFi DNA Assembly Master Mix (E2621, NEB) according to the manufacturer’s instructions. For guide RNA expression targeting NAP1, pX459 was cloned as previously described. Briefly, guides targeting N-terminal of NAP1 were designed using the CHOPCHOP website (https://chopchop.cbu.uib.no/). Oligonucleotides (CAC CGA ACA GTT GTC ATG GAT GCA C, AAA CGT GCA TCC ATG ACA ACT GTT) from IDT were annealed and inserted into the pX459 plasmid after BbsI (NEB) digestion. To develop the FKBP12^F36V^ degradation tag-NAP1 DLD-1 cell, DLD-1 cells were cotransfected using Lipofectamin 3000 (Invitrogen) using above donor vector and pX459 plasmid targeting NAP1. Cells were selected by puromycin (1mg/ml) and diluted into 96 well plates to select for single colony clones. gDNA was extracted from individual clones using lysis buffer (25mM KCl, 5mM Tris HCl pH 8.0, 1.25mM MgCl2, 0.2% NP40, 0.2% Tween-20, 0.4 ug/ml Proteinase K) and heated at 65 °C for 10 min and 98 °C for 5 min incubation. PCR for genotyping was conducted using the following primers: AAG AAC TTT TGA AAA TTT ATA AAT TGA G, GAA AAA TAT TTG GAA TAT AAC TCC AAG.

### Cell synchronization

Nocodazole treatment: Cells were incubated with 1ug/ml nocodazole (Sigma) containing medium to synchronize at the G2/M border for 16 hours and collected for further experiments. R0-3306 treatment: To synchronize at G2, cells were reversibly incubated with 91M RO-3306 (TCI America) containing medium for 20 hrs and collected at the following time points corresponding to their respective cell cycle stages when released in normal growth medium. 0hr – G2; 1hr – M (metaphase); 7hrs – G1. Mitotic shake was employed to obtain maximum number of mitotic cells.

### Cell collection and treatment for Phospho-proteomics

WT HeLa and two independent TBK1 CRISPR knockout lines (TBK1 KO clone 2 and clone 4) were treated with 9uM RO-3306 to synchronize at G2. Cells were released in normal growth medium for an hour to collect approximately 20M mitotic cells per sample (using mitotic shake). To account for off-target gene editing effect, additional groups were added with TBK1 inhibitor treatment. For this, WT HeLa and both TBK1 KO clones were also treated 9μm with R0-3306 for 19.5 hours. 30 mins prior to the G2 wash, the cells were treated with 3μm MRT67307 along with 9μm R03306. After G2 wash the cells were released in normal growth medium with 3μm MRT67307 for 1 hour to collect approximately 20M mitotic cells per sample. 4 biological replicates were collected for each group (Figure 1A).

### Proteomics - cell lysis and protein digestion

At the indicated times, cells were washed twice with ice cold PBS and snap frozen. Cell pellets were lysed in lysis buffer (25 mM EPPS pH 8.5, 8 M Urea, 150 mM NaCl, phosphatase and protease inhibitor cocktail (in-house)), to produce whole cell extracts. Whole cell extracts were sonicated and clarified by centrifugation (16000×g for 10 min at 4°C) and protein concentrations determined by the Bradford assay. Protein extracts (3 mg) were subjected to disulfide bond reduction with 5 mM TCEP (room temperature, 10 min) and alkylation with 25 mM chloroacetamide (room temperature, 20 min). Methanol–chloroform precipitation was performed prior to protease digestion. In brief, four parts of neat methanol were added to each sample and vortexed, one part chloroform was then added to the sample and vortexed, and finally three parts water was added to the sample and vortexed. The sample was centrifuged at 6 000 rpm for 5 min at room temperature and subsequently washed twice with 100% methanol. Samples were resuspended in 100 mM EPPS pH8.5 containing 6 M Urea and digested at 37°C for 2h with Lys-C at a 100:1 protein-to-protease ratio. Samples were then diluted to 0.5 M Urea with 100 mM EPPS pH8.5 solution, trypsin was then added at a 100:1 protein-to-protease ratio and the reaction was incubated for 6 h at 37 °C. Digestion efficiency of a small aliquot was tested, Samples were acidified with 0.1% Trifluoroacetic acid (TFA) final and subjected to C18 solid-phase extraction (SPE) (Sep-Pak, Waters).

### Proteomics - Fe2+-NTA phosphopeptide enrichment

Phosphopeptides were enriched using Pierce High-Select Fe2+-NTA phosphopeptide enrichment kit (Thermo Fisher Scientific, A32992) following the provided protocol. In brief, dried peptides were enriched for phosphopeptides and eluted into a tube containing 25 μL 10% formic acid (FA) to neutralize the pH of the elution buffer and dried down. The unbound peptides (flow through) and washes were combined and saved for total proteome analysis.

### Proteomics - tandem mass tag labeling

Proline-based reporter isobaric Tandem Mass Tag (TMTpro) labeling of dried peptide samples resuspended in 100 mM EPPS pH 8.5, was carried out as followed. For total proteome analysis (50 mg of flow through peptide) and for phosphopeptide proteomics, (desalted, eluted peptides from phospho-enrichment step), 10 μL of a 12.5 μg/μL stock of TMTpro reagent was added to samples, along with acetonitrile to achieve a final acetonitrile concentration of approximately 30% (v/v). Following incubation at room temperature for 1 h, labeling efficiency of a small aliquot was tested for each set (total proteome and phospho-proteome), and the reaction was then quenched with hydroxylamine to a final concentration of 0.5% (v/v) for 15 min. The TMTpro-labeled samples were pooled together at a 1:1 ratio. The total proteome sample and phospho-proteome sample were vacuum centrifuged to near dryness and subjected to C18 solid-phase extraction (SPE) (50 mg, Sep-Pak, Waters).

### Proteomics - off-line basic pH reversed-phase (BPRP) fractionation

Dried TMTpro-labeled sample was resuspended in 100 μl of 10 mM NH4HCO3 pH 8.0 and fractionated using basic pH reverse phase HPLC [79]. Briefly, samples were offline fractionated over a 90 min run, into 96 fractions by high pH reverse-phase HPLC (Agilent LC1260) through an 1) aeris peptide xb-c18 column (Phenomenex; 250 mm x 3.6 mm) for total proteome, 2) kinetex EVO-c18 column (Phenomenex; 150 mm x 2.1 mm) for phosphor-proteome, with mobile phase A containing 5% acetonitrile and 10 mM NH4HCO3 in LC-MS grade H2O, and mobile phase B containing 90% acetonitrile and 10 mM NH4HCO3 in LC-MS grade H2O (both pH 8.0). The 96 resulting fractions were then pooled in a non-continuous manner into 24 fractions (as outlined in Supplemental Figure 5 of [80]) used for subsequent mass spectrometry analysis. Fractions were vacuum centrifuged to near dryness. Each consolidated fraction was desalted via StageTip, dried again via vacuum centrifugation, and reconstituted in 5% acetonitrile, 1% formic acid for LC-MS/MS processing.

### Proteomics – total proteomics analysis using TMTpro

Mass spectrometry data were collected using an Orbitrap Eclipse Tribrid mass spectrometer (Thermo Fisher Scientific, San Jose, CA) coupled to an UltiMate 3000 RSLCnano system liquid chromatography (LC) pump (Thermo Fisher Scientific). Peptides were separated on a 100 μm inner diameter microcapillary column packed in house with ∼40 cm of HALO Peptide ES-C18 resin (2.7 µm, 160 Å, Advanced Materials Technology, Wilmington, DE) with a gradient consisting of 5%–21% (0-85 min), 21-28% (85-110min) (ACN, 0.1% FA) over a total 120 min run at ∼500 nL/min. For analysis, we loaded 1/10 of each fraction onto the column. Each analysis used the Multi-Notch MS^3^-based TMT method [81], to reduce ion interference compared to MS^2^ quantification [82], combined with the FAIMS Pro Interface (using previously optimized 3 CV parameters for TMT multiplexed samples [83] and combined with newly implemented Real Time Search analysis software [84, 85]. The scan sequence began with an MS^1^ spectrum (Orbitrap analysis; resolution 120,000 at 200 Th; mass range 400−1500 m/z; automatic gain control (AGC) target 4×10^5^; maximum injection time 50 ms). Precursors for MS^2^ analysis were selected using a cycle type of 1.25 sec/CV method (FAIMS CV=-40/-60/-80). MS^2^ analysis consisted of collision-induced dissociation (quadrupole ion trap analysis; Rapid scan rate; AGC 1.0×10^4^; isolation window 0.5 Th; normalized collision energy (NCE) 35; maximum injection time 35 ms). Monoisotopic peak assignment was used, and previously interrogated precursors were excluded using a dynamic window (180 s ±10 ppm). Following acquisition of each MS^2^ spectrum, a synchronous-precursor-selection (SPS) API-MS^3^ scan was collected on the top 10 most intense ions b or y-ions matched by the online search algorithm in the associated MS^2^ spectrum [84, 85]. MS^3^ precursors were fragmented by high energy collision-induced dissociation (HCD) and analyzed using the Orbitrap (NCE 45; AGC 2.5×10^5^; maximum injection time 200 ms, resolution was 50,000 at 200 Th). The closeout was set at two peptides per protein per fraction, so that MS^3^s were no longer collected for proteins having two peptide-spectrum matches (PSMs) that passed quality filters [85].

### Proteomics – phosphoproteomics analysis using TMTpro

Mass spectrometry data were collected using an Orbitrap Eclipse Tribrid mass spectrometer (Thermo Fisher Scientific, San Jose, CA) coupled to an UltiMate 3000 RSLCnano system liquid chromatography (LC) pump (Thermo Fisher Scientific). Peptides were separated on a 50 cm µPAC column (PharmaFluidics, Ghent, Belgium) with a gradient consisting of 3%–18% (0-85 min), 18-25% (85-110min) (ACN, 0.1% FA) over a total 125 min run at ∼250 nL/min. For analysis, we loaded half of each fraction onto the column. Each analysis used the FAIMS Pro Interface (using previously optimized 3 CV parameters for TMTpro-labeled phosphopeptides [86]) to reduce ion interference. The scan sequence began with an MS^1^ spectrum (Orbitrap analysis; resolution 120,000 at 200 Th; mass range 400−1500 m/z; automatic gain control (AGC) target 4×10^5^; maximum injection time 50 ms). Precursors for MS^2^ analysis were selected using a cycle type of 1.25 sec/CV method (FAIMS CV=-40/-60/-80). MS^2^ analysis consisted of high energy collision-induced dissociation (HCD) (Orbitrap analysis; resolution 50,000 at 200 Th; isolation window 0.5 Th; normalized collision energy (NCE) 38; AGC 2×10^5^; maximum injection time 86 ms). Monoisotopic peak assignment was used and previously interrogated precursors were excluded using a dynamic window (120 s ±10 ppm).

### Proteomics – data analysis

Mass spectra were processed using a Comet-based (2020.01 rev. 4) software pipeline (Eng et al., 2013). Spectra were converted to mzXML and monoisotopic peaks were re-assigned using Monocle [87]. MS/MS spectra were matched with peptide sequences using the Comet algorithm [88] along with a composite sequence database including the Human Reference Proteome (2020-01 - SwissProt entries only) UniProt database, as well as sequences of common contaminants. This database was concatenated with one composed of all protein sequences in the reversed order. Searches were performed using a 50 ppm precursor ion tolerance for analysis. For total proteomic analysis, the recommended product ion parameters for ion trap ms/ms were used (1.0005 tolerance, 0.4 offset (mono masses), theoretical fragment ions = 1). For phospho proteomics analysis, the recommended product ion parameters for high resolution ms/ms were used (0.02 tolerance, 0.0 offset (mono masses), theoretical fragment ions = 1). TMTpro tags on lysine residues and peptide N termini (+304.207 Da) and carbamidomethylation of cysteine residues (+57.021 Da) were set as static modifications, while oxidation of methionine residues (+15.995 Da) was set as a variable modification. For phosphorylation dataset search, phosphorylation (+79.966 Da) on Serine or Threonine were set as additional variable modifications. Peptide-spectrum matches (PSMs) were adjusted to a 1% false discovery rate (FDR) [89]. PSM filtering was performed using a linear discriminant analysis, [90], while considering the following parameters: Comet Log Expect, Diff Seq. Delta Log Expect, missed cleavages, peptide length, charge state, and precursor mass accuracy. For protein-level comparisons, PSMs were identified, quantified, and collapsed to a 1% peptide false discovery rate (FDR) and then collapsed further to a final protein-level FDR of 1%[91]. Moreover, protein assembly was guided by principles of parsimony to produce the smallest set of proteins necessary to account for all observed peptides. For TMTpro-based reporter ion quantitation, we extracted the summed signal-to-noise (S:N) ratio for each TMTpro channel and found the closest matching centroid to the expected mass of the TMT reporter ion (integration tolerance of 0.003 Da). Reporter ion intensities were adjusted to correct for the isotopic impurities of the different TMTpro reagents according to manufacturer specifications. Proteins were quantified by summing reporter ion signal-to-noise measurements across all matching PSMs, yielding a ‘‘summed signal-to-noise’’ measurement. For total proteome, PSMs with poor quality, MS^3^ spectra with 8 or more TMTpro reporter ion channels missing, or isolation specificity less than 0.7, or with TMT reporter summed signal-to-noise ratio that were less than 160 or had no MS^3^ spectra were excluded from quantification. For phospho proteome, PSMs with poor quality, MS^3^ spectra with 12 or more TMT reporter ion channels missing, or isolation specificity less than 0.8, or with TMT reporter summed signal-to-noise ratio that were less than 160 or had no MS^3^ spectra were excluded from quantification. Phosphorylation site localization was determined using the AScorePro algorithm [92, 93]. AScore is a probability-based approach for high-throughput protein phosphorylation site localization. Specifically, a threshold of 13 corresponded to 95% confidence in site localization.

Protein or peptide quantification values were exported for further analysis in Microsoft Excel, GraphPad Prism, R package and Perseus [94]. Each reporter ion channel was summed across all quantified proteins and normalized assuming equal protein loading of all samples. Phospho peptides were normalized to the protein abundance value (when available) and then normalization of dataset was performed using PhosR package [95]. Gene Ontology enrichment analyses were performed with R package ClusterProfiler (4.0) [96].

Supplemental Data Tables list all quantified proteins as well as associated TMT reporter ratio to control channels used for quantitative analysis.

### Serine/threonine kinase predictions

Kinase predictions were based on experimental biochemical data of their substrate motifs. We had utilized synthetic peptide libraries, containing 198 peptide mixtures, that explored amino acid preference up to 5 residues N-terminal and C-terminal to the phosphorylated Ser/Thr to determine the optimal substrate sequence specificity for recombinant Ser/Thr kinases. In total, 303 kinases were profiled. Their motifs were quantified into position specific scoring matrices (PSSMs) and then applied computationally to score phosphorylation sites based on their surrounding amino acid sequences. These PSSMs were ranked against each site to identify the most favorable kinases. This work is in preparation [58].

### The Kinase Library enrichment analysis

The phosphorylation sites detected in this study were scored by all the characterized kinase PSSMs (303 S/T kinases), and their ranks were determined [58]. For every non-duplicate, singly phosphorylated site, kinases that ranked within the top-15 out of the 303 S/T total kinases were considered as biochemically predicted kinases for their respective phosphorylation site. For assessing kinase motif enrichment, we compared the percentage of phosphorylation sites for which each kinase was predicted among the downregulated/upregulated phosphorylation sites (sites with |log_2_ fold change| greater than or equal 1 and with FDR less than or equal to 0.1), versus the percentage of biochemically favored phosphorylation sites for that kinase within the set of unregulated sites in this study (sites with |log_2_ fold change| less than 1 and with FDR greater than 0.1). Statistical significance was determined using one-sided Fisher’s exact test, and the corresponding p-values were adjusted using the Benjamini-Hochberg procedure. Kinases that were significant (adjusted *p*-value 0.1) for both upregulated and downregulated analysis were excluded from downstream analysis. Then, for every kinase, the most significant enrichment side (upregulated or downregulated) was selected based on the adjusted *p*-value and presented in the scatterplot.

### Western blots

For immunoblotting, cells were lysed using 1X RIPA buffer (Thermo scientific Pierce ™ RIPA Buffer) containing 1X protease/phosphatase inhibitor cocktails (Thermo Scientific Halt™). Protein concentration was quantified using DC™ Protein Assay Kit (Bio-Rad). Human tissue samples were obtained using the INSTA-Blot Human Tissues pre-run western blot (Novus Biologicals). Cell lysates were boiled for 15 mins with 2X LDS buffer containing 50 mM DTT, and 20ug of protein lysates were resolved by 4%-12% Bis-Tris gels and transferred to PVDF membranes. Blots were blocked using 5% non-fat powdered milk in 1X TBST (150mM NaCl, 20mM Tris, pH 8.0, 0.1% Tween 20). Primary and secondary antibody incubations were carried out in 2.5% non-fat powdered milk in 1X TBST for overnight at 4°C and 1hr at room temperature, respectively. Blots were exposed using Clarity™ Western ECL Substrates (Bio-Rad), ECL™ Select Western Blotting Detection Reagent (GE Healthcare), or SuperSignal™ West Femto Maximum Sensitivity Substrate (Thermo Scientific) and detected by the ChemiDoc Imaging System (BioRad).

### PhosTag gels

For protein phosphorylation analysis, 6% Supersep Phos-tag™ gels (Wako Chemicals 192-17401) were used. These gels were run in 1% Tris Glycine and SDS buffer for approximately 2 hours. To increase the transfer efficiency the, phostage gels were soaked for 10 mins (3 times each) in the general transfer buffer containing 5mM EDTA. After washing off the EDTA from the gels using normal transfer buffer, conventional western blot transfer, blocking and antibody incubation steps were followed. For lysates that underwent phosphatase treatment, cells were lysed using 1X RIPA buffer (Thermo scientific Pierce ™ RIPA Buffer) containing 1X protease inhibitor cocktail-EDTA free (Sigma). Approximately 200-250mg of protein lysate was treated with 5U of FastAP alkaline phosphatase (ThermoFisher) for 1 hr at 37°C.

### Immunocytochemistry

Cells for immuno-fluorescence imaging were plated in 6 well cell culture plates (Corning Incorporated) on glass coverslips. Cells were fixed with 4% PFA for 10 mins and permeabilized with 0.1% Triton-X-100 for 10 mins followed by blocking with 10% BSA 5% NGS for 45 mins at RT. Cells were incubated with primary antibodies (diluted in 5% BSA and 2.5% NGS) overnight at 4ºC followed by washing with 1X PBS and incubated with AlexaFluor (Thermo Fisher) conjugated secondary antibodies in the dark for 1 hour. Following the washing step, the cells were stained with .11g/mL DAPI for 5 mins (Thermo Fisher) and mounted on the slides using Fluoromount (Southern-Biotech). Imaging was carried out using a Nikon C2 confocal microscope. For p-TBK1 centrosomal intensity analysis, a different permeabilization method was performed. Cells were permeabilized with 100% ice cold methanol in the refrigerator for 10 mins. For mitotic index, and multinucleated cell count, and cytokinetic cell count random fields of view were captured for each genotype to sample approximately 1000 cells for each biological replicate (n=3). Mitotic cells were identified by chromosome condensation, kinetochore staining by CREST and verified by α-tubulin morphology. For mitotic defects analysis random fields of view were captured for each genotype to sample approximately 50 mitotic cells per biological replicate, and for cytokinetic defects analysis, random fields of view were captured for each genotype to sample approximately 30 cytokinetic cells per biological replicate.

### Cell viability assay for growth curve

Approximately, 200 to 400 cells were plated (4 wells/genotype) in white-coated 96-well plates (Brand Tech Scientific) in growth media. Cell growth curve was obtained by CellTiter-Glo® Luminescent Cell Viability Assay (Promgea) using a luminescence reader every 24 hours. Mean cell number corresponding to the luminescence on each day was normalized to the first day in the graph.

### Immunoprecipitation

Cells were lysed with the following lysis buffer: 50 mM Tris/HCl pH 7.5, 150 mM NaCl, 1 mM EGTA, 1 mM EDTA, 0.5(v/v) NP-40, 1 mM sodium orthovanadate, 50 mM NaF, 5 mM sodium pyrophosphate, 0.27 M sucrose, 10 mM Na 2-glycerophosphate, 0.2 mM phenylmethylsulphonyl fluoride, 1x protease/phosphatase inhibitor cocktail (Pierce), and 50 mM iodoacetamide. Cells were incubated for 20 minutes end over end at 4°C then spun at 16,000 x *g* for 15 minutes at 4°C. Supernatant was measured using the Dc™ protein assay (Bio-Rad). 500 ug of protein was incubated on magnetic beads (FLAG M2 beads (Sigma), GFP beads (Chromotek)) end over end at 4°C for 2 hrs. For endogenous IPs, supernatant was precleared on TrueBlot Protein G magnetic beads (Rockland) for 30 min at 4°C. 1.2 mg of protein was incubated with 2uL of TBK1 antibody for 1hr at 4°C (#3013S, Cell Signaling). Magnetic beads were added for 1hr at 4°C. 2X LDS with 50mM DTT was used to elute protein off the beads, and pH was restored with NaOH.

### Transfection

HEK293T cells were plated on poly-d-lysine coated dishes and reverse transfected with 1:1 jetOPTIMUS transfection reagent (Polyplus). Cells were treated the next day with 100ng/ml nocodazole (Sigma) and collected 16hrs later. HeLa cells were reverse transfected with 1:3 or 1:6 XtremeGENE 9™ (Roche) transfection reagent according to the manufacturer’s instructions.

### qPCR

RNA was isolated using Trizol Reagent (Ambion Life Technologies) and converted to cDNA using iScript™ (Bio-Rad) per the manufacturer’s directions. SYBR Green Supermix (Bio-Rad), 20 ng of cDNA and 0.4 μM of each primer set was mixed in a 10 μl RT-PCR reaction that was ran on the CFX96 System (Bio-Rad). The primers that were used span exons: Actin F: 5’ CCCGCCGCCAGCTCACCAT 3’, R: 5’ CGATGGAGGGGAAGACGGCCC 3’. TANK F: 5’ AGCAAGGAGTCTTGGCAGTC 3’, R: 5’ GCACTGTGTTTCAGTTGCAGT 3’. SINTBAD F: 5’ ACCAGTTCCAGCATGAGTTACA 3’, R: 5’ TCTCCCTCAGCTCTGTCTCC 3’. AZI2 F: 5’ AGGTGGAAACTCAGCAGGTG 3’, R: 5’ ATGGATCCGTTTGTTTGGCT 3’. IL-6 F: 5’ AGCCACTCACCTCCTCAGAACGAA 3’, R: 5’ AGTGCCTCTTTGCTGCTTTCACAC 3’. TNF-α F: 5’ TCAATCGGCCCGACTATCTC 3’, R: 5’ CAGGGCAATGATCCCAAAGT 3’. IL-10 F: 5’ AAGACCCAGACATCAAGGCG 3’, R: 5’ CAGGGAAGAAATCGATGACAGC 3’. RT-PCR was performed in triplicate wells from three independent biological experiments. Expression levels were normalized to β-actin and fold change was determined by comparative C_T_ method.

### Statistical analysis

For comparisons between two groups, student’s t-test was used to determine statistical significance. Ordinary one-way ANOVA followed by Tukey’s multiple comparisons were used for three or more groups using GraphPad Prism software. Additional details are available in the figure legends. Differences in means were considered significant if p <0.05 and designated as the following p<0.05 - *; p< 0.01 - **; p< 0.001 - ***. p<.0001 - ****; ns – not significant.

## Acknowledgments

We thank Dr. Daniela Cimini and Mathew Bloomfield for advice, reagents, and their expertise. The authors declare no competing financial interests.

## Author contributions

Conceptualization: AMP

Planning and methodology: SP, SAS, AO, AMP

Experimentation: SP, SAS, KN, LZ, SRB, LEF, ND, AO, AMP

Data Analysis: SP, SAS, KN, TMY, JMJ, EMH

Reagents: SAS

Computational Software for Analysis: LCC

Writing – original draft: SP, AMP

Writing – review &editing: SP, SAS, AO, AMP

All authors have read and approved the manuscript.

## Data and materials availability

MS data will be submitted to MassIVE Repository. Constructs used for this study will be available through Addegene.org. Raw western blotting images will be deposited on Mendeley Data. All other reagents, data, and material requests will be fulfilled by the corresponding author, Alicia M. Pickrell, Ph.D. All data are available in the main text or the supplementary materials.

## References

1. Nasa, I. and A.N. Kettenbach, Coordination of Protein Kinase and Phosphoprotein Phosphatase Activities in Mitosis. Front Cell Dev Biol, 2018. 6: p. 30.

2. Nigg, E.A., Mitotic kinases as regulators of cell division and its checkpoints. Nat Rev Mol Cell Biol, 2001. 2(1): p. 21–32.

3. Seki, A., et al., Bora and the kinase Aurora a cooperatively activate the kinase Plk1 and control mitotic entry. Science, 2008. 320(5883): p. 1655–8.

4. Enserink, J.M. and R.D. Kolodner, An overview of Cdk1-controlled targets and processes. Cell Div, 2010. 5: p. 11.

5. Dephoure, N., et al., A quantitative atlas of mitotic phosphorylation. Proc Natl Acad Sci U S A, 2008. 105(31): p. 10762–7.

6. Olsen, J.V., et al., Quantitative phosphoproteomics reveals widespread full phosphorylation site occupancy during mitosis. Sci Signal, 2010. 3(104): p. ra3.

7. Colas, P., Cyclin-dependent kinases and rare developmental disorders. Orphanet J Rare Dis, 2020. 15(1): p. 203.

8. Schneider, I. and J. Ellenberg, Mysteries in embryonic development: How can errors arise so frequently at the beginning of mammalian life? PLoS Biol, 2019. 17(3): p. e3000173.

9. Huang, K.L., et al., Spatially interacting phosphorylation sites and mutations in cancer. Nat Commun, 2021. 12(1): p. 2313.

10. Singh, V., et al., Phosphorylation: Implications in Cancer. Protein J, 2017. 36(1): p. 1–6.

11. Pillai, S., et al., Tank binding kinase 1 is a centrosome-associated kinase necessary for microtubule dynamics and mitosis. Nat Commun, 2015. 6: p. 10072.

12. Sarraf, S.A., et al., PINK1/Parkin Influences Cell Cycle by Sequestering TBK1 at Damaged Mitochondria, Inhibiting Mitosis. Cell Rep, 2019. 29(1): p. 225–235 e5.

13. Chen, W., et al., TBK1 Promote Bladder Cancer Cell Proliferation and Migration via Akt Signaling. J Cancer, 2017. 8(10): p. 1892–1899.

14. Uhlen, M., et al., A pathology atlas of the human cancer transcriptome. Science, 2017. 357(6352).

15. Wei, C., et al., Elevated expression of TANK-binding kinase 1 enhances tamoxifen resistance in breast cancer. Proc Natl Acad Sci U S A, 2014. 111(5): p. E601–10.

16. Bonnard, M., et al., Deficiency of T2K leads to apoptotic liver degeneration and impaired NF-kappaB-dependent gene transcription. EMBO J, 2000. 19(18): p. 4976–85.

17. Maan, M., et al., Tank Binding Kinase 1 modulates spindle assembly checkpoint components to regulate mitosis in breast and lung cancer cells. Biochim Biophys Acta Mol Cell Res, 2021. 1868(3): p. 118929.

18. Onorati, M., et al., Zika Virus Disrupts Phospho-TBK1 Localization and Mitosis in Human Neuroepithelial Stem Cells and Radial Glia. Cell Rep, 2016. 16(10): p. 2576–2592.

19. Thurston, T.L., et al., Recruitment of TBK1 to cytosol-invading Salmonella induces WIPI2-dependent antibacterial autophagy. EMBO J, 2016. 35(16): p. 1779–92.

20. Goncalves, A., et al., Functional dissection of the TBK1 molecular network. PLoS One, 2011. 6(9): p. e23971.

21. Heo, J.M., et al., The PINK1-PARKIN Mitochondrial Ubiquitylation Pathway Drives a Program of OPTN/NDP52 Recruitment and TBK1 Activation to Promote Mitophagy. Mol Cell, 2015. 60(1): p. 7–20.

22. Clark, K., et al., The TRAF-associated protein TANK facilitates cross-talk within the IkappaB kinase family during Toll-like receptor signaling. Proc Natl Acad Sci U S A, 2011. 108(41): p. 17093–8.

23. Gatot, J.S., et al., Lipopolysaccharide-mediated interferon regulatory factor activation involves TBK1-IKKepsilon-dependent Lys(63)-linked polyubiquitination and phosphorylation of TANK/I-TRAF. J Biol Chem, 2007. 282(43): p. 31131–46.

24. Tanaka, Y. and Z.J. Chen, STING specifies IRF3 phosphorylation by TBK1 in the cytosolic DNA signaling pathway. Sci Signal, 2012. 5(214): p. ra20.

25. Bakshi, S., et al., Identification of TBK1 complexes required for the phosphorylation of IRF3 and the production of interferon beta. Biochem J, 2017. 474(7): p. 1163–1174.

26. Fujita, F., et al., Identification of NAP1, a regulatory subunit of IkappaB kinase-related kinases that potentiates NF-kappaB signaling. Mol Cell Biol, 2003. 23(21): p. 7780–93.

27. Zhang, C., et al., Structural basis of STING binding with and phosphorylation by TBK1. Nature, 2019. 567(7748): p. 394–398.

28. Ryzhakov, G. and F. Randow, SINTBAD, a novel component of innate antiviral immunity, shares a TBK1-binding domain with NAP1 and TANK. EMBO J, 2007. 26(13): p. 3180–90.

29. Gleason, C.E., et al., Polyubiquitin binding to optineurin is required for optimal activation of TANK-binding kinase 1 and production of interferon beta. J Biol Chem, 2011. 286(41): p. 35663–35674.

30. Lazarou, M., et al., The ubiquitin kinase PINK1 recruits autophagy receptors to induce mitophagy. Nature, 2015. 524(7565): p. 309–314.

31. Moore, A.S. and E.L. Holzbaur, Dynamic recruitment and activation of ALS-associated TBK1 with its target optineurin are required for efficient mitophagy. Proc Natl Acad Sci U S A, 2016. 113(24): p. E3349–58.

32. Richter, B., et al., Phosphorylation of OPTN by TBK1 enhances its binding to Ub chains and promotes selective autophagy of damaged mitochondria. Proc Natl Acad Sci U S A, 2016. 113(15): p. 4039–44.

33. Kim, J.Y., et al., Dissection of TBK1 signaling via phosphoproteomics in lung cancer cells. Proc Natl Acad Sci U S A, 2013. 110(30): p. 12414–9.

34. Sasai, M., et al., Cutting Edge: NF-kappaB-activating kinase-associated protein 1 participates in TLR3/Toll-IL-1 homology domain-containing adapter molecule-1-mediated IFN regulatory factor 3 activation. J Immunol, 2005. 174(1): p. 27–30.

35. Sasai, M., et al., NAK-associated protein 1 participates in both the TLR3 and the cytoplasmic pathways in type I IFN induction. J Immunol, 2006. 177(12): p. 8676–83.

36. Fu, T., et al., Mechanistic insights into the interactions of NAP1 with the SKICH domains of NDP52 and TAX1BP1. Proc Natl Acad Sci U S A, 2018. 115(50): p. E11651–E11660.

37. Larabi, A., et al., Crystal structure and mechanism of activation of TANK-binding kinase 1. Cell Rep, 2013. 3(3): p. 734–46.

38. Ma, X., et al., Molecular basis of Tank-binding kinase 1 activation by transautophosphorylation. Proc Natl Acad Sci U S A, 2012. 109(24): p. 9378–83.

39. Li, F., et al., Structural insights into the interaction and disease mechanism of neurodegenerative disease-associated optineurin and TBK1 proteins. Nat Commun, 2016. 7: p. 12708.

40. Thurston, T.L., et al., The TBK1 adaptor and autophagy receptor NDP52 restricts the proliferation of ubiquitin-coated bacteria. Nat Immunol, 2009. 10(11): p. 1215–21.

41. Morton, S., et al., Enhanced binding of TBK1 by an optineurin mutant that causes a familial form of primary open angle glaucoma. FEBS Lett, 2008. 582(6): p. 997–1002.

42. Ravenhill, B.J., et al., The Cargo Receptor NDP52 Initiates Selective Autophagy by Recruiting the ULK Complex to Cytosol-Invading Bacteria. Mol Cell, 2019. 74(2): p. 320–329 e6.

43. Wild, P., et al., Phosphorylation of the autophagy receptor optineurin restricts Salmonella growth. Science, 2011. 333(6039): p. 228–33.

44. Vargas, J.N.S., et al., Spatiotemporal Control of ULK1 Activation by NDP52 and TBK1 during Selective Autophagy. Mol Cell, 2019. 74(2): p. 347–362 e6.

45. Wong, Y.C. and E.L. Holzbaur, Optineurin is an autophagy receptor for damaged mitochondria in parkin-mediated mitophagy that is disrupted by an ALS-linked mutation. Proc Natl Acad Sci U S A, 2014. 111(42): p. E4439–48.

46. Pourcelot, M., et al., The Golgi apparatus acts as a platform for TBK1 activation after viral RNA sensing. BMC Biol, 2016. 14: p. 69.

47. Nabet, B., et al., The dTAG system for immediate and target-specific protein degradation. Nat Chem Biol, 2018. 14(5): p. 431–441.

48. Nabet, B., et al., Rapid and direct control of target protein levels with VHL-recruiting dTAG molecules. Nat Commun, 2020. 11(1): p. 4687.

49. Fagerberg, L., et al., Analysis of the human tissue-specific expression by genome-wide integration of transcriptomics and antibody-based proteomics. Mol Cell Proteomics, 2014. 13(2): p. 397–406.

50. Fitzgerald, K.A., et al., IKKepsilon and TBK1 are essential components of the IRF3 signaling pathway. Nat Immunol, 2003. 4(5): p. 491–6.

51. Sharma, S., et al., Triggering the interferon antiviral response through an IKK-related pathway. Science, 2003. 300(5622): p. 1148–51.

52. Tojima, Y., et al., NAK is an IkappaB kinase-activating kinase. Nature, 2000. 404(6779): p. 778–82.

53. Taylor, J.H., Nucleic acid synthesis in relation to the cell division cycle. Ann N Y Acad Sci, 1960. 90: p. 409–21.

54. Prescott, D.M. and M.A. Bender, Synthesis of RNA and protein during mitosis in mammalian tissue culture cells. Exp Cell Res, 1962. 26: p. 260–8.

55. Hyer, M.L., et al., A small-molecule inhibitor of the ubiquitin activating enzyme for cancer treatment. Nat Med, 2018. 24(2): p. 186–193.

56. Clark, K., et al., Novel cross-talk within the IKK family controls innate immunity. Biochem J, 2011. 434(1): p. 93–104.

57. Li, J., et al., TMTpro reagents: a set of isobaric labeling mass tags enables simultaneous proteome-wide measurements across 16 samples. Nat Methods, 2020. 17(4): p. 399–404.

58. Johnson, J.L., et al., A global atlas of substrate specificities for the human serine/threonine kinome. bioRxiv, 2022: p. 2022.05.22.492882.

59. Hirota, T., et al., Histone H3 serine 10 phosphorylation by Aurora B causes HP1 dissociation from heterochromatin. Nature, 2005. 438(7071): p. 1176–80.

60. Steigemann, P., et al., Aurora B-mediated abscission checkpoint protects against tetraploidization. Cell, 2009. 136(3): p. 473–84.

61. Ozlu, N., et al., Binding partner switching on microtubules and aurora-B in the mitosis to cytokinesis transition. Mol Cell Proteomics, 2010. 9(2): p. 336–50.

62. Carmena, M., S. Ruchaud, and W.C. Earnshaw, Making the Auroras glow: regulation of Aurora A and B kinase function by interacting proteins. Curr Opin Cell Biol, 2009. 21(6): p. 796–805.

63. Bitan, A., et al., Asymmetric microtubule function is an essential requirement for polarized organization of the Drosophila bristle. Mol Cell Biol, 2010. 30(2): p. 496–507.

64. Shapiro, R.S. and K.V. Anderson, Drosophila Ik2, a member of the I kappa B kinase family, is required for mRNA localization during oogenesis. Development, 2006. 133(8): p. 1467–75.

65. Abdu, U., D. Bar, and T. Schupbach, spn-F encodes a novel protein that affects oocyte patterning and bristle morphology in Drosophila. Development, 2006. 133(8): p. 1477–84.

66. Dubin-Bar, D., et al., The Drosophila IKK-related kinase (Ik2) and Spindle-F proteins are part of a complex that regulates cytoskeleton organization during oogenesis. BMC Cell Biol, 2008. 9: p. 51.

67. Lin, T., et al., Spindle-F Is the Central Mediator of Ik2 Kinase-Dependent Dendrite Pruning in Drosophila Sensory Neurons. PLoS Genet, 2015. 11(11): p. e1005642.

68. Barros, C.S. and T. Bossing, Microtubule disruption upon CNS damage triggers mitotic entry via TNF signaling activation. Cell Rep, 2021. 36(1): p. 109325.

69. Kodani, A., et al., Zika virus alters centrosome organization to suppress the innate immune response. EMBO Rep, 2022. 23(9): p. e52211.

70. Fukasaka, M., et al., Critical role of AZI2 in GM-CSF-induced dendritic cell differentiation. J Immunol, 2013. 190(11): p. 5702–11.

71. Barbie, D.A., et al., Systematic RNA interference reveals that oncogenic KRAS-driven cancers require TBK1. Nature, 2009. 462(7269): p. 108–12.

72. Zhu, L., et al., TBKBP1 and TBK1 form a growth factor signalling axis mediating immunosuppression and tumourigenesis. Nat Cell Biol, 2019. 21(12): p. 1604–1614.

73. Katz, M., I. Amit, and Y. Yarden, Regulation of MAPKs by growth factors and receptor tyrosine kinases. Biochim Biophys Acta, 2007. 1773(8): p. 1161–76.

74. Pylayeva-Gupta, Y., E. Grabocka, and D. Bar-Sagi, RAS oncogenes: weaving a tumorigenic web. Nat Rev Cancer, 2011. 11(11): p. 761–74.

75. Yamamoto, T., et al., Continuous ERK activation downregulates antiproliferative genes throughout G1 phase to allow cell-cycle progression. Curr Biol, 2006. 16(12): p. 1171–82.

76. Meloche, S., Cell cycle reentry of mammalian fibroblasts is accompanied by the sustained activation of p44mapk and p42mapk isoforms in the G1 phase and their inactivation at the G1/S transition. J Cell Physiol, 1995. 163(3): p. 577–88.

77. Jones, S.M. and A. Kazlauskas, Growth-factor-dependent mitogenesis requires two distinct phases of signalling. Nat Cell Biol, 2001. 3(2): p. 165–72.

78. Damhofer, H., A. Radzisheuskaya, and K. Helin, Generation of locus-specific degradable tag knock-ins in mouse and human cell lines. STAR Protoc, 2021. 2(2): p. 100575.

79. Wang, Y., et al., Reversed-phase chromatography with multiple fraction concatenation strategy for proteome profiling of human MCF10A cells. Proteomics, 2011. 11(10): p. 2019–26.

80. Paulo, J.A., et al., Quantitative mass spectrometry-based multiplexing compares the abundance of 5000 S. cerevisiae proteins across 10 carbon sources. J Proteomics, 2016. 148: p. 85–93.

81. McAlister, G.C., et al., MultiNotch MS3 enables accurate, sensitive, and multiplexed detection of differential expression across cancer cell line proteomes. Anal Chem, 2014. 86(14): p. 7150–8.

82. Paulo, J.A., J.D. O’Connell, and S.P. Gygi, A Triple Knockout (TKO) Proteomics Standard for Diagnosing Ion Interference in Isobaric Labeling Experiments. J Am Soc Mass Spectrom, 2016. 27(10): p. 1620–5.

83. Schweppe, D.K., et al., Characterization and Optimization of Multiplexed Quantitative Analyses Using High-Field Asymmetric-Waveform Ion Mobility Mass Spectrometry. Anal Chem, 2019. 91(6): p. 4010–4016.

84. Erickson, B.K., et al., Active Instrument Engagement Combined with a Real-Time Database Search for Improved Performance of Sample Multiplexing Workflows. J Proteome Res, 2019. 18(3): p. 1299–1306.

85. Schweppe, D.K., et al., Full-Featured, Real-Time Database Searching Platform Enables Fast and Accurate Multiplexed Quantitative Proteomics. J Proteome Res, 2020. 19(5): p. 2026–2034.

86. Schweppe, D.K., et al., Optimized Workflow for Multiplexed Phosphorylation Analysis of TMT-Labeled Peptides Using High-Field Asymmetric Waveform Ion Mobility Spectrometry. J Proteome Res, 2020. 19(1): p. 554–560.

87. Rad, R., et al., Improved Monoisotopic Mass Estimation for Deeper Proteome Coverage. J Proteome Res, 2021. 20(1): p. 591–598.

88. Eng, J.K., T.A. Jahan, and M.R. Hoopmann, Comet: an open-source MS/MS sequence database search tool. Proteomics, 2013. 13(1): p. 22–4.

89. Elias, J.E. and S.P. Gygi, Target-decoy search strategy for increased confidence in large-scale protein identifications by mass spectrometry. Nat Methods, 2007. 4(3): p. 207–14.

90. Huttlin, E.L., et al., A tissue-specific atlas of mouse protein phosphorylation and expression. Cell, 2010. 143(7): p. 1174–89.

91. Savitski, M.M., et al., A Scalable Approach for Protein False Discovery Rate Estimation in Large Proteomic Data Sets. Mol Cell Proteomics, 2015. 14(9): p. 2394–404.

92. Beausoleil, S.A., et al., A probability-based approach for high-throughput protein phosphorylation analysis and site localization. Nat Biotechnol, 2006. 24(10): p. 1285–92.

93. Gassaway, B.M., et al., A multi-purpose, regenerable, proteome-scale, human phosphoserine resource for phosphoproteomics. Nat Methods, 2022.

94. Tyanova, S., et al., The Perseus computational platform for comprehensive analysis of (prote)omics data. Nat Methods, 2016. 13(9): p. 731–40.

95. Kim, H.J., et al., PhosR enables processing and functional analysis of phosphoproteomic data. Cell Rep, 2021. 34(8): p. 108771.

96. Wu, T., et al., clusterProfiler 4.0: A universal enrichment tool for interpreting omics data. Innovation (N Y), 2021. 2(3): p. 100141.

