## Supplemental figures for "NAK associated protein 1/NAP1 is required for mitosis and cytokinesis by activating TBK1"

Paul *et al.*

* Alicia M. Pickrell.

**This PDF file includes:**

Figs. S1 to S5

**Other Supplementary Materials for this manuscript include the following:**
Data S1 to S4

Figure S1

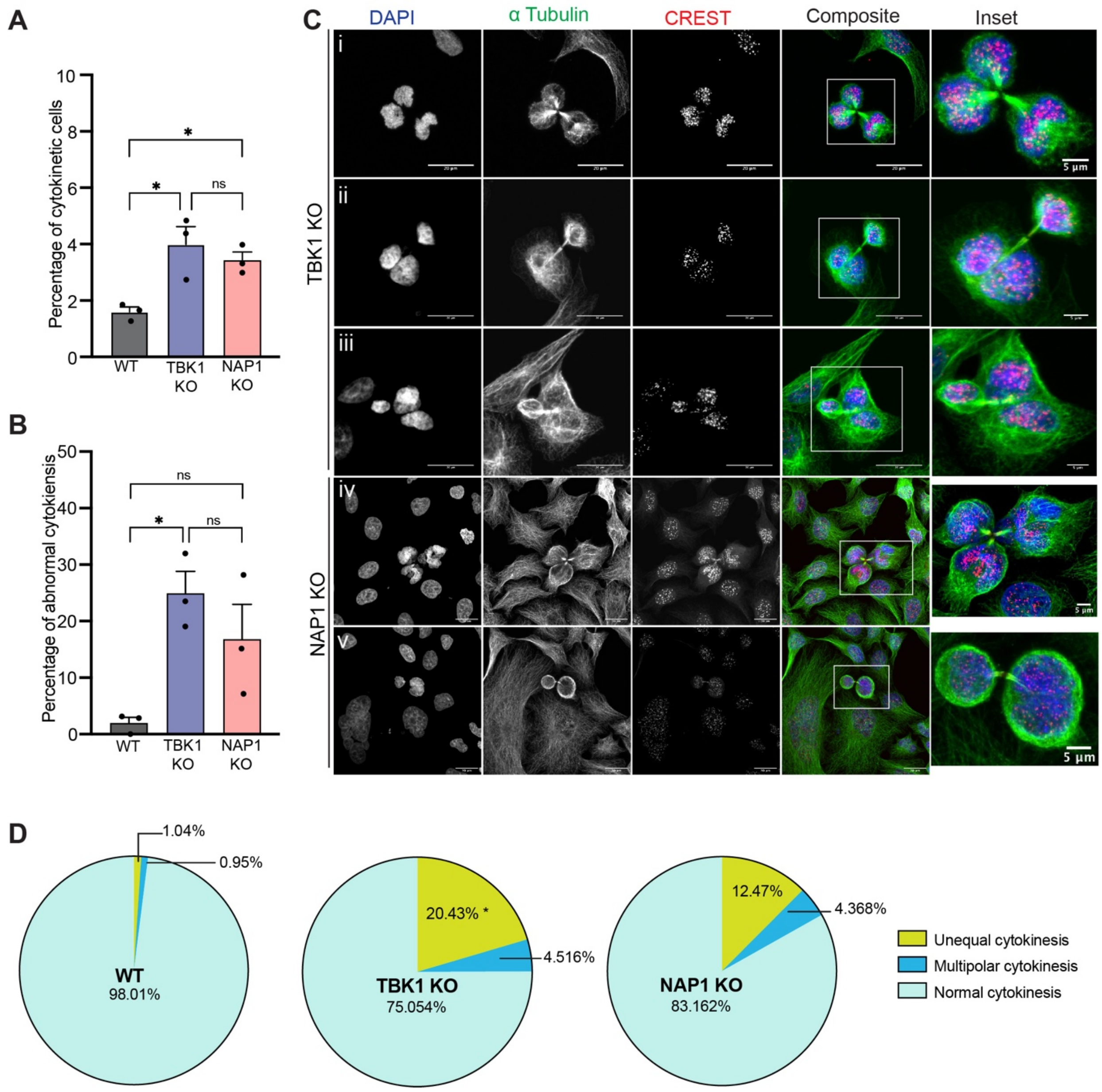


**Fig. S2: Loss of NAP1 and TBK1 cause similar cytokinetic defects.**

**(A-B)** Cytokinetic (A) and abnormal cytokinetic (B) cell count percentages from an asynchronous population of WT HeLa, TBK1 KO and NAP1 KO. Error bars indicate ±SD; n=3 independent experiments. Random fields of view were captured sampling approximately 1000 cells per biological replicate from each genotype.

**(C)** Representative confocal images of cytokinetic defects seen in TBK1 KO (i-iii) and NAP1 KO (iv-v): (i) multipolar cytokinesis, (ii) unequal cytokinesis, (iii) combination of unequal multipolar cytokinesis, (iv) multipolar cytokinesis, (v) unequal cytokinesis. DAPI (blue) was used as a nuclear counterstain, α-tubulin for cytoskeleton staining (green), and CREST for kinetochore staining (red). Scale bar, 20 μm, insets, 5μm.

(**D**) Pie chart representing the percentage of different types of cytokinetic defects found in HeLa, TBK1, and NAP1 KO cells. Random fields of view were captured sampling approximately 30-40 cytokinetic cells per biological replicate from each genotype. * $p < .05$ compared to HeLa.

One way ANOVA was performed for all statistical analysis. * $p < .05$, ns = not significant.

**Figure S2**

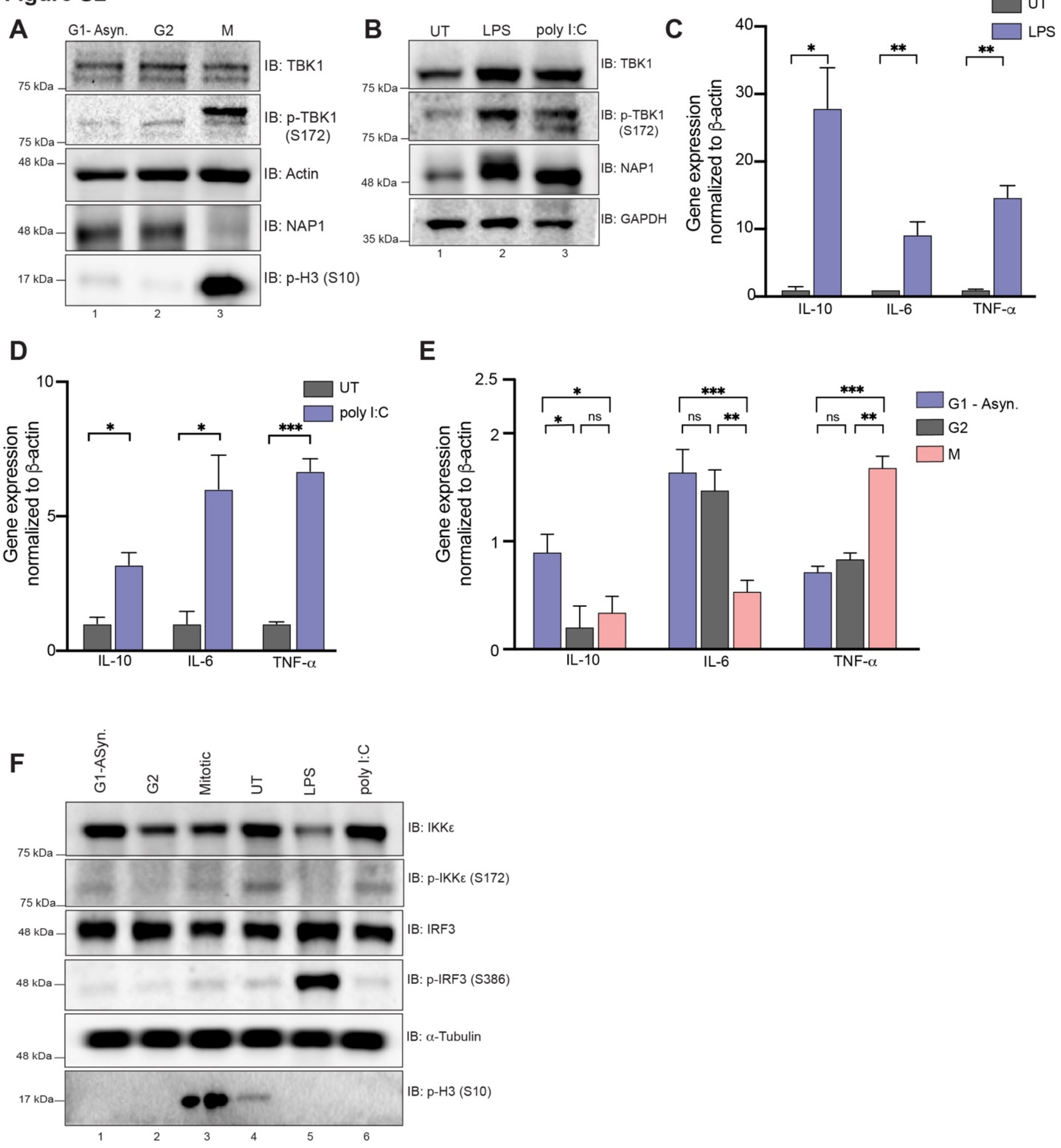


**Fig. S2: Mitosis does not elicit an innate immune response.**

**(A)** Western blot analysis of THP-1 cells synchronized at G2, M, or G1-Async to determine p-TBK1 levels. Cells were synchronized at G2 using RO-3306. G2 cells were released for approximately 45-60 minutes to collect mitotic cells and for approximately 7 hours to collect G1 samples.

(**B)** Western blot analysis of THP-1 cells stimulated with LPS or poly I:C for 1 hrs and 8 hrs, respectively. Blots were probed for p-TBK1 and NAP1. Cells were synchronized using RO-3306.
(**C-D)** Relative mRNA expression of cytokines upregulated during innate immunity normalized to □-actin when stimulated with LPS (C) for 1 hrs or poly I:C (D) for 8 hrs. Error bars indicate ±SD; n = 3 independent experiments.
(**E)** Relative mRNA expression of cytokines during different cell cycle stages normalized to □-actin. Error bars indicate ±SD; n = 3 independent experiments. Cells were synchronized at G2 using RO-3306. G2 cells were released for approximately 45-60 minutes to collect mitotic and for approximately 7 hours to collect G1 samples.
(**F)** Western blot analysis of p-IRF3 and p-IKKe of THP-1 cells synchronized at G2, M, or G1-Async or stimulated with LPS or poly I:C for 1 hrs and 8 hrs, respectively. Cells were synchronized at G2 using RO-3306. G2 cells were released for approximately 45-60 minutes to collect mitotic, and for approximately 7 hours to collect G1 samples.
Student's t-test or one way ANOVA was performed for all statistical analysis. * $p < .05$, ** $p < .01$, *** $p <.001$, ns = not significant.

Figure S3

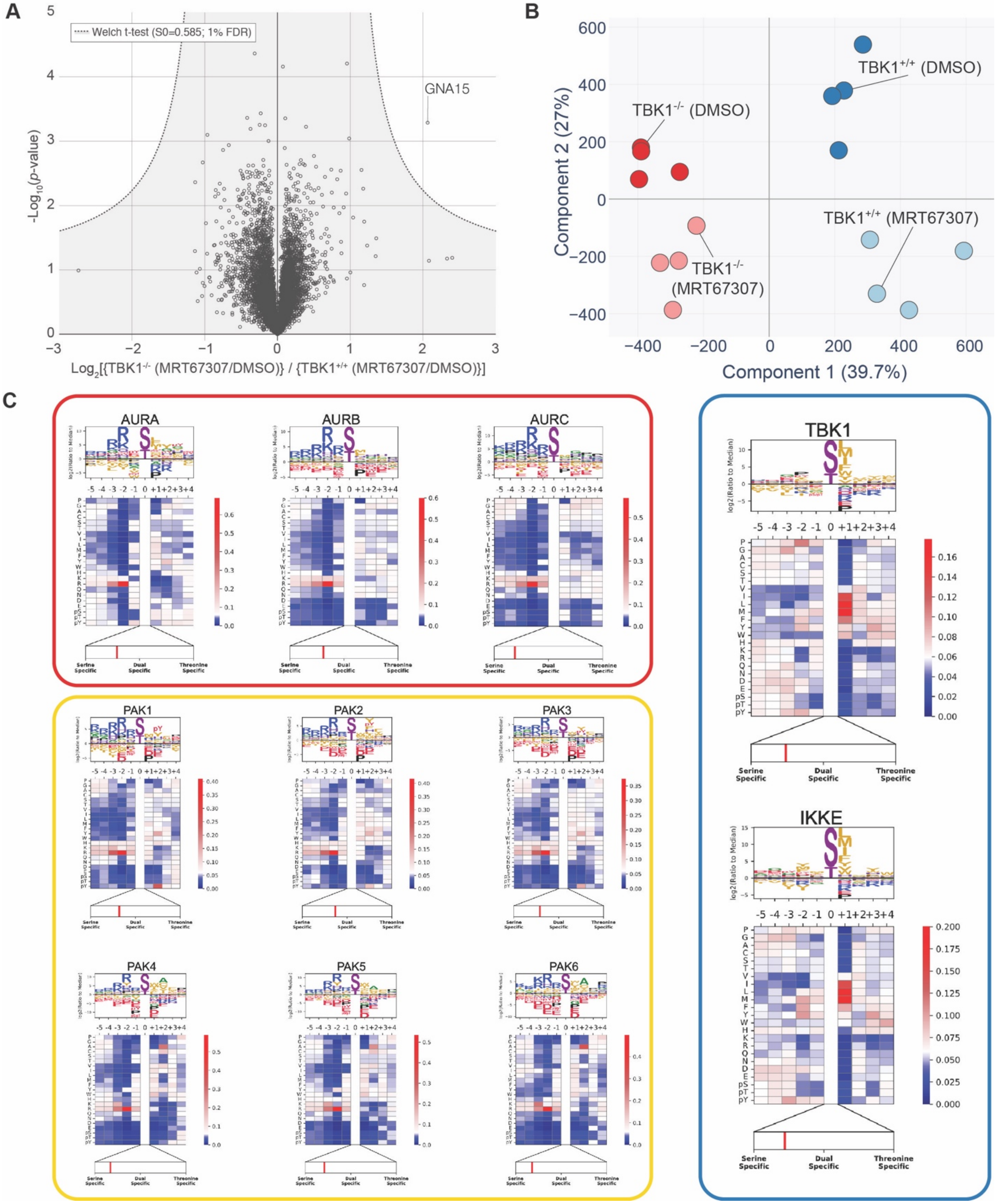


**Fig. S3: Quantitative proteomics pipeline identifies downstream substrates regulated upon mitosis.**

**(A)** Volcano plots [Log10 (p-value) versus Log2 ratio] representing protein abundance that are affected by MRT67307 and loss of TBK1. Proteins are shown in black circles. The inset indicated additional color coding for the statistical analysis.

**(B)** Principal component analysis (PCA) of the phosphoproteome data. Replicate samples are shown separately. 39.7% of the changes in phospho abundance are provided by Component 1, which represents the genetic background component, while 27% of the change are provided by component 2, which represents the small molecule inhibitor treatment.

**(C)** Individual motifs for experimentally-derived substrate sequence specificity for Aurora family, PAK famliy, TBK1 and IKKe kinases. Data derived from (Johnson et al., 2022).

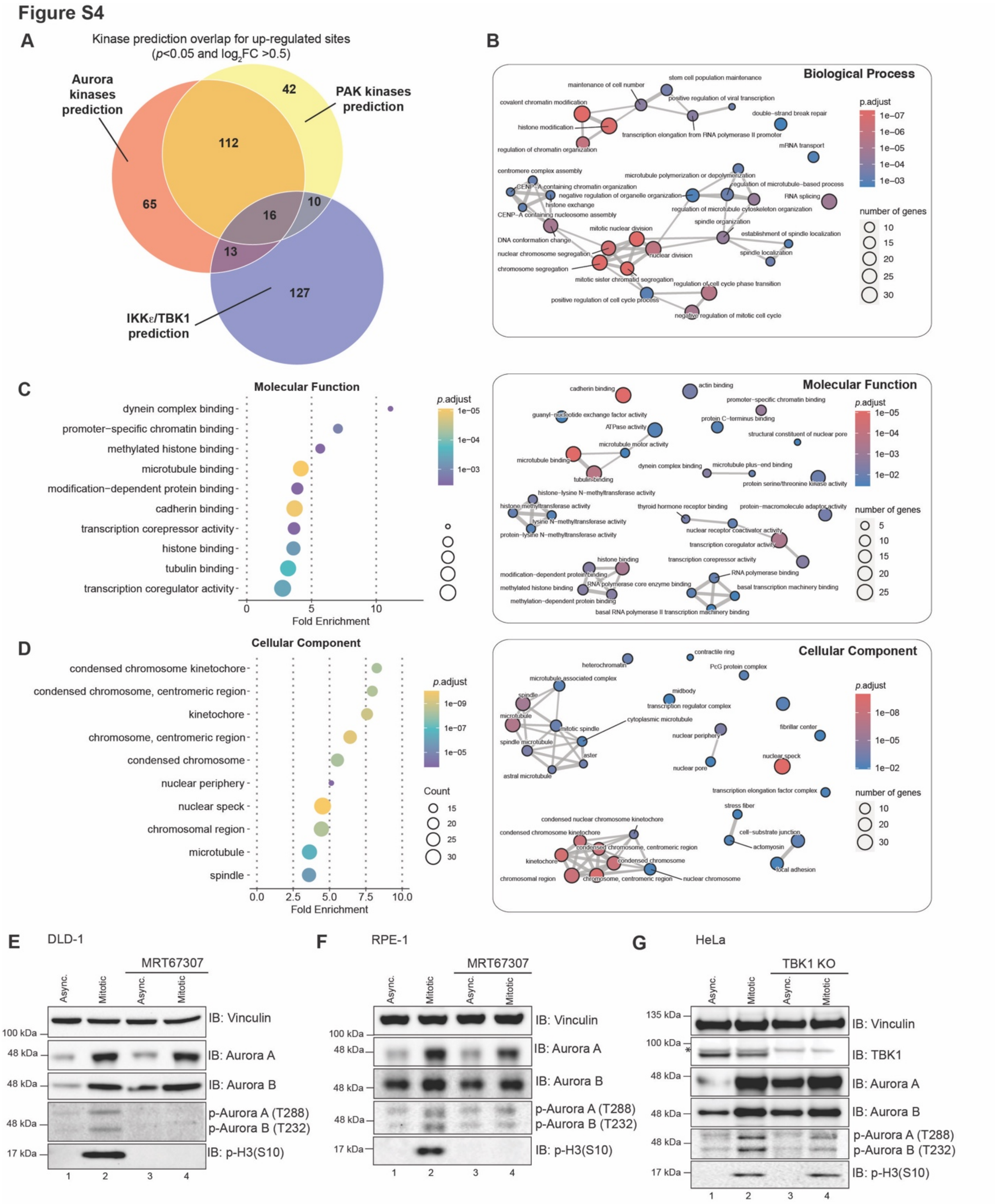


**Figure S7: Quantitative proteomics pipeline identifies TBK1 downstream substrates upon mitosis.**

**(A)** Venn diagram illustrating the kinase prediction overlap for up-regulated sites passing the *p*-value cutoff of $p<0.05$ and $log_2$ ratio cutoff of +0.5. Related to Table S2.

**(B-D)** Gene Ontology terms enrichment analysis and associated enrichment map networks for enriched phosphor-sites (Tier 2). For enrichment map networks, each node represents a gene set (i.e., a GO term) and each edge represents the overlap between two gene sets.

**(E-G)** Western blot analysis of p-Aurora A (T288) and p-Aurora B (T232) in asynchronous and synchronized mitotic cells from DLD-1 (A) and RPE-1 (B) treated with MRT67307 or HeLa WT compared to TBK1 KO cells (C). Cells were synchronized at G2 using RO-3306 prior to mitotic release. MRT67307 treatment occurred 1 hr prior to G2 and throughout release. Asynchronous cells were treated for 2 hrs.

Figure S5

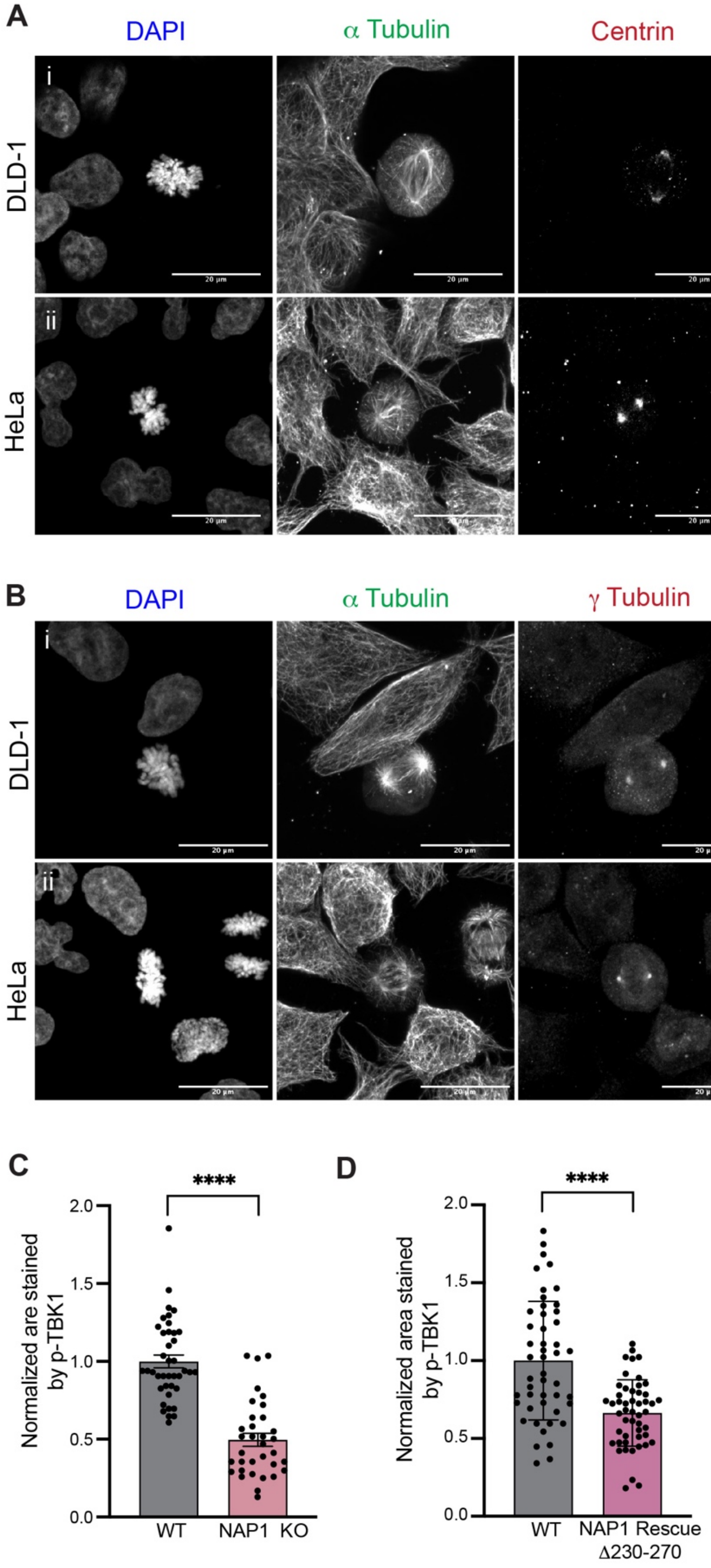


**Figure S5: Activated TBK1 is not confined to only centrosomes.**

(**A**) Representative confocal images of Centrin (red) and p-TBK1 (magenta) staining in (i) DLD-1, (ii) HeLa cells. DAPI (blue) was used as a nuclear counterstain, and $\alpha$-tubulin for cytoskeleton staining (green). Scale bar, 20 μm.

(**B**) Representative confocal images of g-tubulin (red) and p-TBK1 (magenta) staining in (i) DLD-1, (ii) HeLa cells. DAPI (blue) was used as a nuclear counterstain, and $\alpha$-tubulin for cytoskeleton staining (green). Scale bar, 20 μm.

**(C)** Relative area of p-TBK1 staining around centrosomes of mitotic cells from WT HeLa and NAP1 KO. 40-50 mitotic cells per group were quantified from 2 biological replicates. Error bars indicate ±SEM.

**(D)** Relative area of p-TBK1 staining around centrosomes of mitotic cells from WT HeLa and stable EGFP NAP1 $\Delta$230-270 rescue cells. 40-50 mitotic cells per group were quantified from 2 biological replicates. Error bars indicate ±SEM.

Student's t-test was performed for all statistical analysis. **** $p < .0001$.

**Data S1. (separate file)**

Relevant to **Fig. 6 b, c, e** and **Supplementary Fig. 3a-d**

**Data S2 (separate file)**

Relevant to **Fig. 6 b, c, e** and **Supplementary Fig. 3a-d**

**Data S3 (separate file)**

Relevant to **Fig. 6 b, c, d** and **Supplementary Fig. 3c, 4a**

**Data S4 (separate file)**

Relevant to **Fig. 7**
